# Permeability of the HIV-1 capsid to metabolites modulates viral DNA synthesis

**DOI:** 10.1101/2020.04.30.071217

**Authors:** Chaoyi Xu, Douglas K. Fischer, Sanela Rankovic, Wen Li, Rob Dick, Brent Runge, Roman Zadorozhnyi, Jinwoo Ahn, Christopher Aiken, Tatyana Polenova, Alan N. Engelman, Zandrea Ambrose, Itay Rousso, Juan R. Perilla

**Author notes:** Contributed equally (Chaoyi Xu, Douglas K. Fischer, Sanela Rankovic).

## Abstract

Reverse transcription, an essential event in the HIV-1 lifecycle, requires deoxynucleotide triphosphates (dNTPs) to fuel DNA synthesis, thus requiring penetration of dNTPs into the viral core. The central cavity of the capsid protein (CA) hexamer reveals itself as a plausible channel that allows the passage of dNTPs into assembled capsids. Nevertheless, the molecular mechanism of nucleotide import into the capsid remains unknown. Employing all-atom molecular dynamics simulations, we established that cooperative binding between nucleotides inside a CA hexamer cavity results in energetically-favorable conditions for passive translocation of dNTPs into the HIV-1 capsid. Furthermore, binding of the host cell metabolite inositol hexakisphosphate (IP_6_) enhances dNTP import, while binding of synthesized molecules like benzenehexacarboxylic acid (BHC) inhibits it. The enhancing effect on reverse transcription by IP_6_ and the consequences of interactions between CA and nucleotides were corroborated using atomic force microscopy, transmission electron microscopy, and virological assays. Collectively, our results provide an atomistic description of the permeability of the HIV-1 capsid to small molecules and reveal a novel mechanism for the involvement of metabolites in HIV-1 capsid stabilization, nucleotide import and reverse transcription.

## Introduction

Fusion between HIV-1 virions and target cells engenders the release of the viral capsid into the host cell cytoplasm. The capsid, a cone-shaped protein assembly composed of ∼250 capsid protein (CA) hexamers and 12 CA pentamers, encapsulates two copies of the viral single-stranded RNA (ssRNA) genome and viral proteins, including reverse transcriptase (RT) and integrase (IN)^1-3^. Successful infection requires reverse transcription of the ssRNA into double-stranded viral DNA (vDNA) and integration of vDNA into the host cell genome. Based on steric hindrance of the intact capsid by the nuclear pore complex^4^, major structural rearrangement or capsid shedding has been postulated to occur prior to or during nuclear import^5,6^, a process commonly referred to as uncoating. While reverse transcription likely initiates within the capsid and is affected by uncoating, the mechanisms connecting the two viral processes remain a long-standing question in HIV-1 biology^7-10^. We and others have proposed that reverse transcription mechanically induces changes in capsid morphology and triggers uncoating^11,12^. To support vDNA synthesis, RT requires adequate concentrations of deoxynucleotide triphosphates (dNTPs) within the lumen of the capsid. The HIV-1 capsid has been described as semipermeable and proposed to regulate the passage of ions including dNTPs from the cytoplasm to its interior^3,4,8^.

Inositol phosphates (IPs) are abundant cellular metabolites that play crucial roles in fundamental cellular processes^13^. Interestingly, ions and small molecules like inositol hexakisphosphate (IP_6_) and benzenehexacarboxylic acid (BHC, or mellitic acid) have been reported to bind to HIV-1 Gag and CA hexamers and impact virion assembly and reverse transcription^4,14-18^. Here, using a multi-pronged approach combining all-atom molecular dynamics (MD) simulations, atomic force microscopy (AFM), transmission electron microscopy (TEM), confocal microscopy, and virus infectivity assays, we have analyzed the effect of IP_6_ on HIV-1 capsid stability and reverse transcription. Furthermore, we determined the molecular mechanism that regulates translocation of dNTPs through the capsid.

## Results and Discussion

The CA hexameric cavity is surrounded by six copies of a β-hairpin, helix 1 and a short loop connecting them (Fig. 1a, b). Its radial profile, illustrated in Fig. 1a for the native CA hexamer^19^, contains an exterior tubular channel and an interior conical volume separated by a 1.4 Å radial constriction near Arg18 residues (Fig. 1a). The inner surface of the outward facing cavity is surrounded by residues Asn5, Gln7, and Gln13 that lie within or nearby the β-hairpin. Helix 1 sequences are nearly identical across primate lentiviruses, including invariant charged residues Arg18, Lys25, Glu28, Glu29, and Lys30 (Fig. 1b). These residues, which account for the electrostatic potential of the hexamer cavity, contrast to the negative charge found in the exposed surfaces of the hexamer (Fig. 1c). Overall, based exclusively on the structural features of CA hexamers, the volume occupied by the ring of six Arg18 side chains represents a steric barrier for translocation of small molecules through the central cavity.

**Figure 1.**
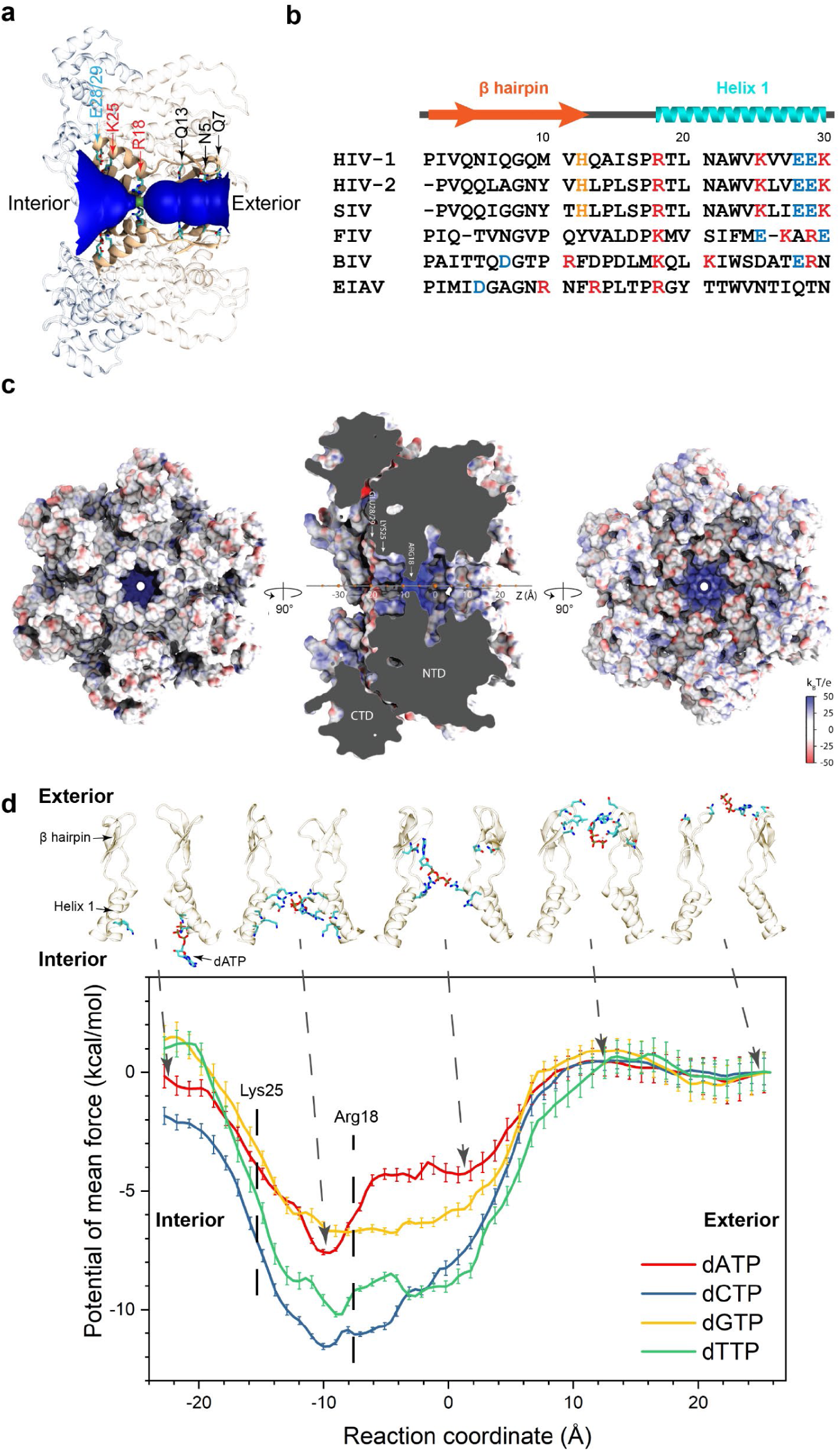
Architecture of the HIV-1 CA hexameric cavity. (a) The radial profile of the native hexamer (PDB accession code 4XFX) showing positions of key CA residues that constitute the inner surface of the central cavity. The interior and exterior of the capsid are located on the left and right of the diagram, respectively. (b) Comparative analysis and sequence conservation of 30 N-terminal CA residues from six lentiviruses (SIV, simian immunodeficiency virus; FIV, feline immunodeficiency virus; BIV, bovine immunodeficiency virus; EIAV, equine infectious anemia virus). Secondary structures are indicated above the sequences. (c) Interior surface, side and exterior surface representing the electrostatic potential. The reaction coordinate (progress variable) for all US/HREX simulations was defined as the position of the center of mass of the nucleotide on the cavity axis. (d) The 1D free energy landscapes of single dNTP translocations through the CA hexamer central cavity. Five representative snapshots are shown indicating the interactions between CA and dNTPs.

To explore the effects of the dynamic behavior of CA hexamers on the permeability of molecules through assembled HIV-1 capsids, we employed free-energy MD simulations to determine the free energy landscapes of dNTPs, IP_6_, and BHC for interacting with the central cavity (Fig. 1d, S1, simulations #1-10). To monitor the displacement of the nucleotide in the cavity, a progress variable (PV) was introduced as the location of the center of mass of each small molecule of interest on the pore axis of the hexamer (Fig. 1c). The resulting free energy landscapes displayed energy basins in proximity to Arg18 for dNTPs, rNTPs, NMPs, IP_6_, and BHC (Fig. 1d, Fig. S1, Table S1). Unexpectedly, the depth of the energy basin differed between nucleotide types, as a minimum free energy of −7 to - 8 kcal/mol was observed for dATP and dGTP, while a minimum of −10 to −12 kcal/mol was observed for dCTP and dTTP. We infer that steric effects of the larger purine nucleotides incur lower binding affinity compared to the more compact pyrimidines. These results reveal nucleoside-specific binding of dNTPs to CA hexamers. However, once bound, release of the dNTP would require overcoming a +barrier of at least 6 kcal/mol; thus, dNTPs are highly unlikely to translocate through CA hexamers on their own.

In our simulations, a single nucleotide becomes trapped within the cavity, as the energy well is too deep to enable its release. Our MD simulations revealed that the central hexamer cavity could accommodate two small molecules simultaneously interacting with the ring of Arg18 residues (movie 1; molecular simulation #25, 26). Therefore, we increased the stoichiometric ratio between small molecules and CA hexamer to 2:1 (simulation #11, #12, #13; Fig. 2, S2) to examine nucleotide translocation in the presence of multiple charged molecules inside the cavity. Specifically, the free energy profile of a first molecule (shown in the horizontal axes of Fig. 2a, c, e) was calculated throughout the cavity in the presence of a second molecule bound to the capsid cavity near Arg18 (shown in the vertical axes of Fig. 2a, c, e). These calculations revealed a second free-energy minimum in the interior of the capsid near Lys25 for a dNTP in the presence of IP_6_ or another dNTP (Movie 2; Fig. 2a, c, S2). Moreover, the minimum action path connecting the interior and exterior of the capsid (dashed lines in Fig. 2a, c, e and Fig. S2) demonstrated a marked difference in the ability of CA hexamers to translocate dNTPs in the presence of BHC, IP_6_ or another dNTP. Specifically, a positive free-energy gradient, which promoted dNTP motion inwards, was observed for dNTPs and IP_6_ (Fig. 2a, 2c, S2b, S2c), while a negative gradient was observed in the presence of BHC (Fig. 2e, S2d). The negative gradient induced by BHC implies that it inhibits nucleotide import. Additionally, the difference in the free energy gradient induced by IP_6_ compared to dNTPs shows that the former is a stronger activator of nucleotide import. Our molecular simulations revealed that IP_6_ will increase dNTP transport, whereas BHC will block or reduce it.

**Figure 2.**
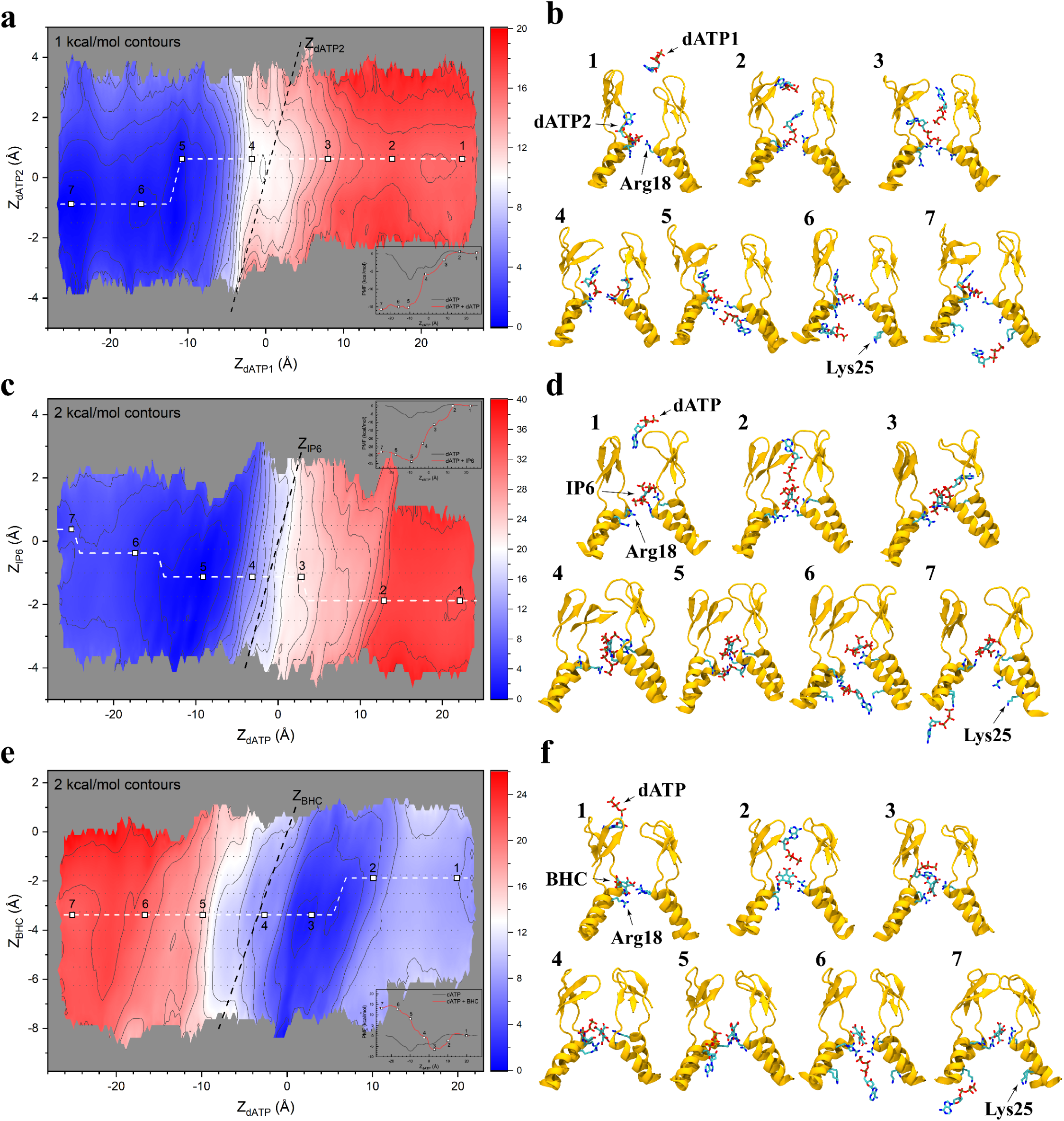
Cooperative binding of small molecules to the central hexamer cavity. Two-dimensional free energy landscapes of dATP translocation through the cavity, in the presence of (a) an additional dATP, (c) IP_6_ or (e) The horizontal reaction coordinates (in Å) represent the location of the center of mass (COM) of a dATP molecule on the cavity axis, which connects the exterior and interior of the capsid (respectively right and left). The vertical reaction coordinates (in Å) are the positions of the COM of (a) a second dATP, (c) IP_6_ or (e) BHC on the progress variable along the central hexamer axis. The magnitudes of the free energies are indicated in the color scalebar in units of kcal/mol. Pathways connecting the interior and exterior are shown as a dashed line and representative structures corresponding to these translocation events are illustrated in (b), (d) and (f).

Alterations of Lys25 were previously found to alter capsid stability but still produce virus-like particles^20,21^. Results from our simulations showed that Lys25 and Glu28 substitutions modified dNTP import (simulations #14-17, Fig. S1d). In particular, K25A was predicted to enhance import while K25E/N were predicted to inhibit translocation, indicating that Lys25 charge is not the sole determinant of nucleotide import. These data indicated that Lys25 plays a role in coordinating solvent molecules around the nucleotide after dewetting of the dNTP through its binding to Arg18 (Fig. S3). Based on these predictions, the effect of K25N on HIV-1 capsid stability and reverse transcription was examined. In vitro assembly reactions assessed by TEM (Fig. S4) and magic angle spinning (MAS) NMR (Fig. S5) revealed that wild-type (WT) and K25N constructs behaved similarly, while K25A constructs were unable to form mature-like conical capsids and were not pursued for further experiments (Fig. S4). Consistent with these observations, K25N HIV-1 was morphologically similar to the WT virus (Fig. 3a). However, K25N capsids were less stable than WT capsids based on CA retention in permeabilized viruses, which was assessed by antibody staining and confocal microscopy (Fig. 3b, S6).

**Figure 3.**
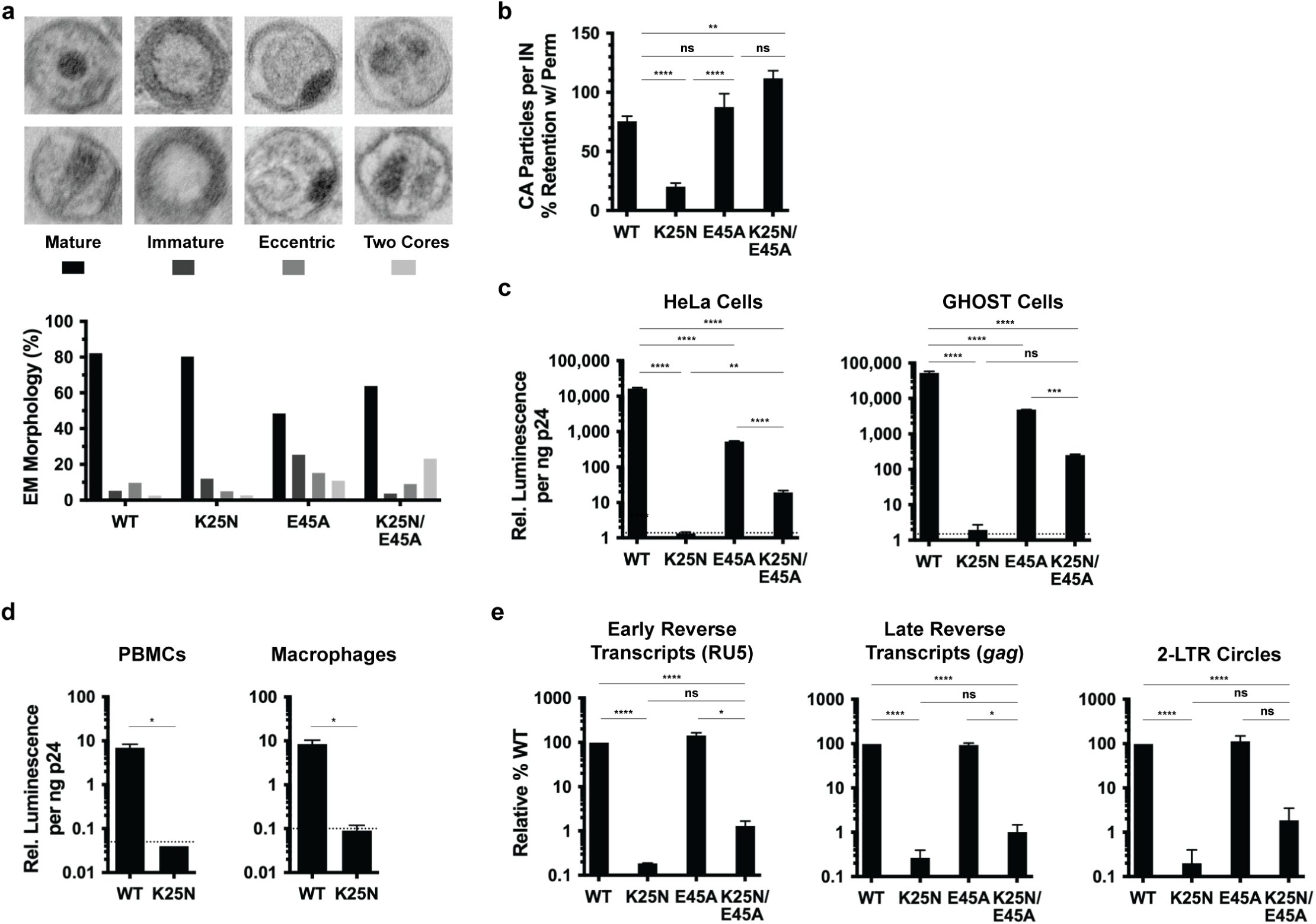
Characterization of K25N HIV-1 and K25N/E45A HIV-1. (a) Transmission electron microscopy (TEM) classification of virus particle morphologies for HIV-1_NL4-3_ bearing indicated CA changes. (b) Viruses bearing the indicated CA substitutions were captured on glass with or without pre-fixation virus membrane permeabilization, followed by immunostaining for CA and imaging. Data are expressed as the percentage of non-permeabilized CA staining retained with membrane permeabilization for each virus. (c, d) Single-cycle infectivities of WT and indicated CA mutant viruses in indicated cells. Dotted lines represent average luciferase signal of uninfected cells. (e) DNA synthesis and 2-LTR circle formation of WT and CA mutant viruses. Error bars indicate SEM for 2-4 experiments. **** P < 0.0001, *** P < 0.001, ** P < 0.01, * P < 0.05.

To evaluate infectivity, cell lines, peripheral blood mononuclear cells (PBMCs) and monocyte-derived macrophages were infected with normalized levels of WT or K25N HIV-1_NL4-3_-based luciferase reporter viruses. While WT HIV-1 infected all cell types tested, K25N was noninfectious (Fig. 3c,d). Similar results were obtained with WT and K25N HIV-1_LAI_ reporter viruses (data not shown). To determine the step of the virus life cycle at which K25N HIV-1 infection was impaired, early and late reverse transcripts and 2-long terminal repeat (LTR)-containing circles were measured by quantitative PCR (qPCR). Levels of K25N early and late reverse transcripts were reduced 540-fold and 385-fold, respectively, relative to WT HIV-1 (Fig. 3e). The number of 2-LTR circles, a surrogate marker of vDNA nuclear entry, was likewise 500-fold lower for K25N compared to WT HIV-1 (Fig. 3e). These data indicate that K25N HIV-1 is defective for reverse transcription.

As K25N HIV-1 capsids were unstable, which itself could result in impaired reverse transcription, we assessed the affect of the capsid hyperstabilizing CA substitution E45A^22,23^ on the stability of K25N capsids. K25N/E45A HIV-1 was more stable than K25N HIV-1 (Fig. 3b), though both E45A HIV-1 and K25N/E45A HIV-1 produced higher percentages of morphologically defective particles than did either WT or K25N HIV-1 (Fig. 3a). E45A HIV-1 was 10- to 30-fold less infectious than WT HIV-1 (Fig. 3c), consistent with prior characterization of this mutant^22^, while K25N/E45A HIV-1 harbored a 200- to 800-fold infectivity defect (Fig. 3c). While E45A HIV-1 synthesized WT levels of early and late reverse transcripts and 2-LTR circles (Fig. 3e), K25N/E45A HIV-1 early and late reverse transcripts were reduced 78-fold and 100-fold, respectively, and 2-LTR circles were reduced 54-fold (Fig. 3e). Thus, the E45A stabilization change can separate the capsid stability and dNTP import phenotypes of K25N HIV-1, suggesting that impaired dNTP import underlies the observed K25N and K25N/E45A HIV-1 reverse transcription and infectivity defects.

Changes in capsid stiffness and morphologies during reverse transcription were monitored in real-time by AFM (Fig. 4)^11,24,25^. The effect of IP_6_ binding on the stiffness of intact HIV-1 capsids was analyzed using AFM operated in the nanoindentation mode. In agreement with our previously reported findings, isolated WT capsid had an average stiffness value of 0.132 ± 0.007 N/m (n=30) (Fig, 4a). Remarkably, binding of IP_6_ to isolated HIV-1 capsids increased their stiffness by nearly 2-fold to an average value of 0.245 ± 0.021 N/m (n=36). Initially, untreated WT capsids exhibited an average stiffness value of 0.131 ± 0.012 N/m; n=11. Upon IP_6_ addition, the value increased to 0.258 ± 0.035 N/m (n=15, time = 0 h), but decreased immediately after reverse transcription was initiated, reaching a minimum of 0.078 ± 0.013 N/m (n=7) at 7 h; the minimum stiffness value was reached earlier for capsids treated with IP_6_ than for capsids without IP_6_ (5 h compared to ∼20 h)^24^. Overall, our AFM measurements show that addition of IP_6_ initially stiffens the capsids, then induces faster softening after initiation of reverse transcription. In contrast, addition of the synthetic molecule BHC resulted in a smaller increase in the stiffness value (0.196 ± 0.043 N/m; n=4; Fig. 4b). The stiffness of BHC-treated capsids remained mostly unchanged during reverse transcription and remained fully intact even after 24 h of reverse transcription. These data are consistent with previous results showing that IP_6_ stabilizes HIV-1 capsids and promotes viral DNA synthesis^14^ and our computational experiments that revealed BHC blocked nucleotide translocation through the CA hexameric cavity.

**Fig 4:**
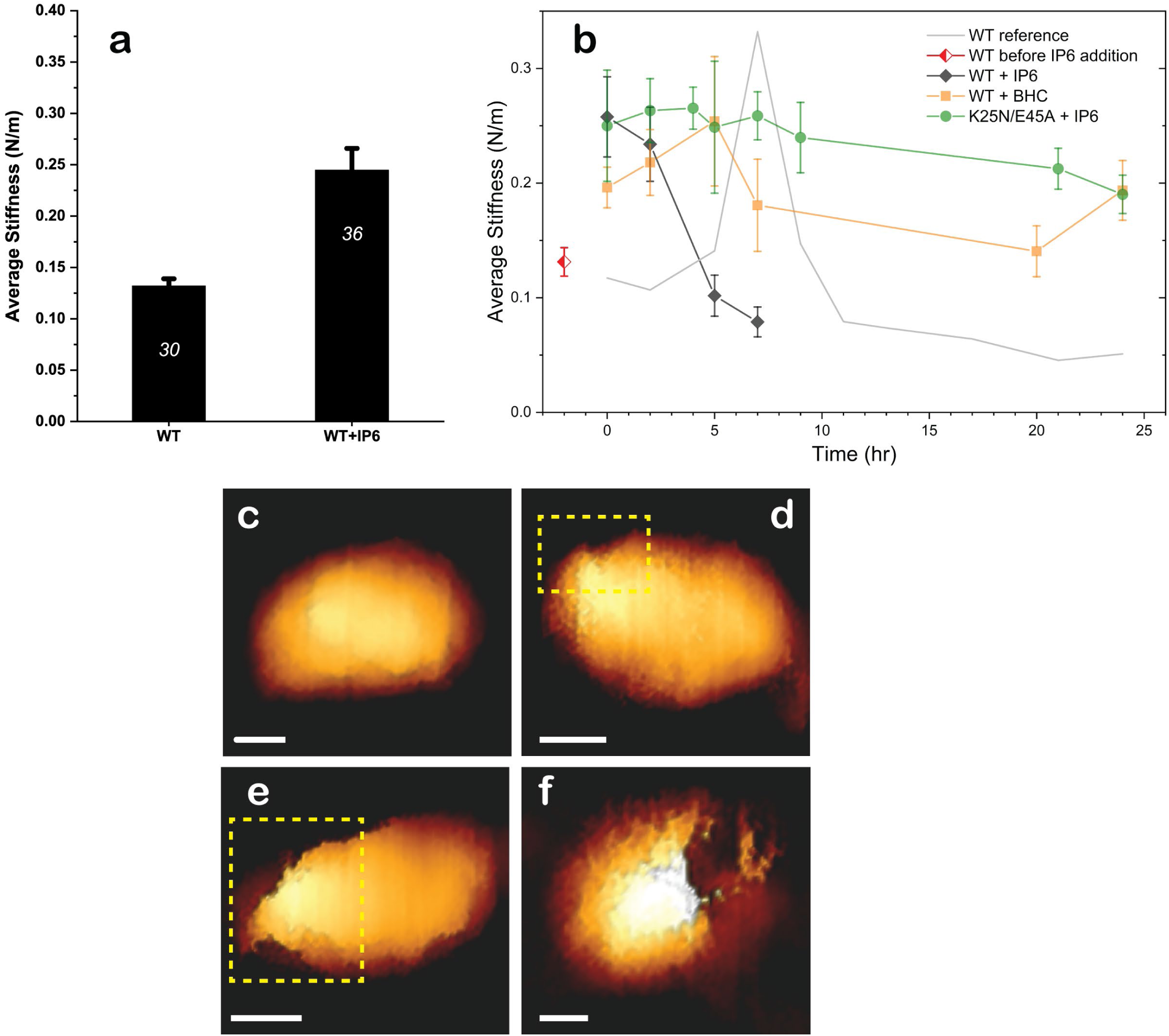
Morphologies and mechanical properties of HIV-1 capsids from AFM. a) The average stiffness of isolated WT HIV −1 capsids in the absence (n=30) and presence (n=36) of IP_6_. Error bars indicate SEM. Statistical significance was confirmed using the Mann-Whitney U test for p < 0.05. (b) Average stiffness of HIV-1 capsids treated with 100 µM IP_6_ (black) or BHC (orange) as a function of time after the addition of dNTPs and MgCl_2_. The red diamond represents the average stiffness of WT capsids before addition of IP_6_ or BHC. The initial average stiffness, at time 0, was measured before addition of dNTPs and MgCl_2_. At each time point, the stiffness of 2 to 4 capsids was measured without stopping the reaction. WT data (taken from reference Rankovic, et al. J Virol 2017) is presented as a gray line for comparison. The error bars represent SEM. (c) A representative cone-shaped capsid (of 20 imaged) prior to addition of dNTP and MgCl_2_. (d-f) Deformed and damaged capsids visualized after 5 h. For clarity, openings in the capsids are shown within a dashed yellow rectangle. Scale bars, 50 nm. A total of 52 capsids were visualized. (Approximately 400 force-distance curves were obtained from individual capsids).

To further characterize the effects of IP_6_ on reverse transcription, we analyzed the morphologies of IP_6_-treated WT capsids during reverse transcription by AFM operated in the quantitative imaging mode. Prior to reverse transcription, capsids had a well-defined conical appearance. A representative capsid (out of a total of 20 that were imaged) is shown in Fig. 4c. After 5 h, openings at various sizes in the capsid appeared (Fig. 4d-f). Similar to our previous findings^26,27^, the openings were localized exclusively at or near the narrow end of the capsids. Untreated and IP_6-_treated HIV-1 capsids underwent complete disassembly during the time course of the experiments. However, complete disassembly was 3.4 times faster in the presence of IP_6_ (7 h vs. to 24 h). Analysis of the reactions beyond 7 h of reverse transcription revealed mostly fragments of various sizes that lacked a defined morphology (from a total of 52 capsids, 11 remained intact). Overall, we find that IP_6_ accelerates capsid disassembly during reverse transcription.

The stiffness of the K25N/E45A isolated capsids was also measured during reverse transcription. While the stability of these capsids was not sufficient to withstand the full duration of the experiment due to spontaneous disassembly, their stiffness remained unchanged over 5 h. Although addition of IP_6_ did not affect CA retention on K25N/E45A capsids (Fig. S7), their stability was increased, which enabled us to measure their stiffness values during 24 h of reverse transcription. In agreement with our virus infectivity and reverse transcription analyses, the stiffness of the double mutant capsids remained unchanged during 24 hours of reverse transcription (Fig 4c), consistent with the observed reverse transcription defect of K25N/E45A HIV-1.

Reverse transcription and the consequent disassembly of the HIV-1 capsid are fueled by the ability of the capsid to import dNTPs from the cytoplasm. Free energy calculations reveal a molecular dynamic view of the interaction between a freely diffusing nucleotide and the HIV-1 CA hexamer (Fig 5). First, a nucleotide freely diffuses from the exterior solvent to the beta-hairpin region. Subsequently, the loss of entropy by binding of the nucleotide to Arg18 is compensated by the strength of the electrostatic interactions between Arg18 and dNTP phosphate residues. Hydrogen bonds between the nucleotide base and polar residues in helix 1 confer distinct binding affinities for individual nucleotide species, which acts as a selectivity filter. However, to be released into the capsid lumen, a single nucleotide needs to overcome a high energetic penalty (free energy barrier of over 6 kcal/mol). Engagement of a second charged molecule such as dNTP or IP_6_ within the channel provides the required thermodynamic effector to release the initial dNTP within the capsid lumen.

**Figure 5.**
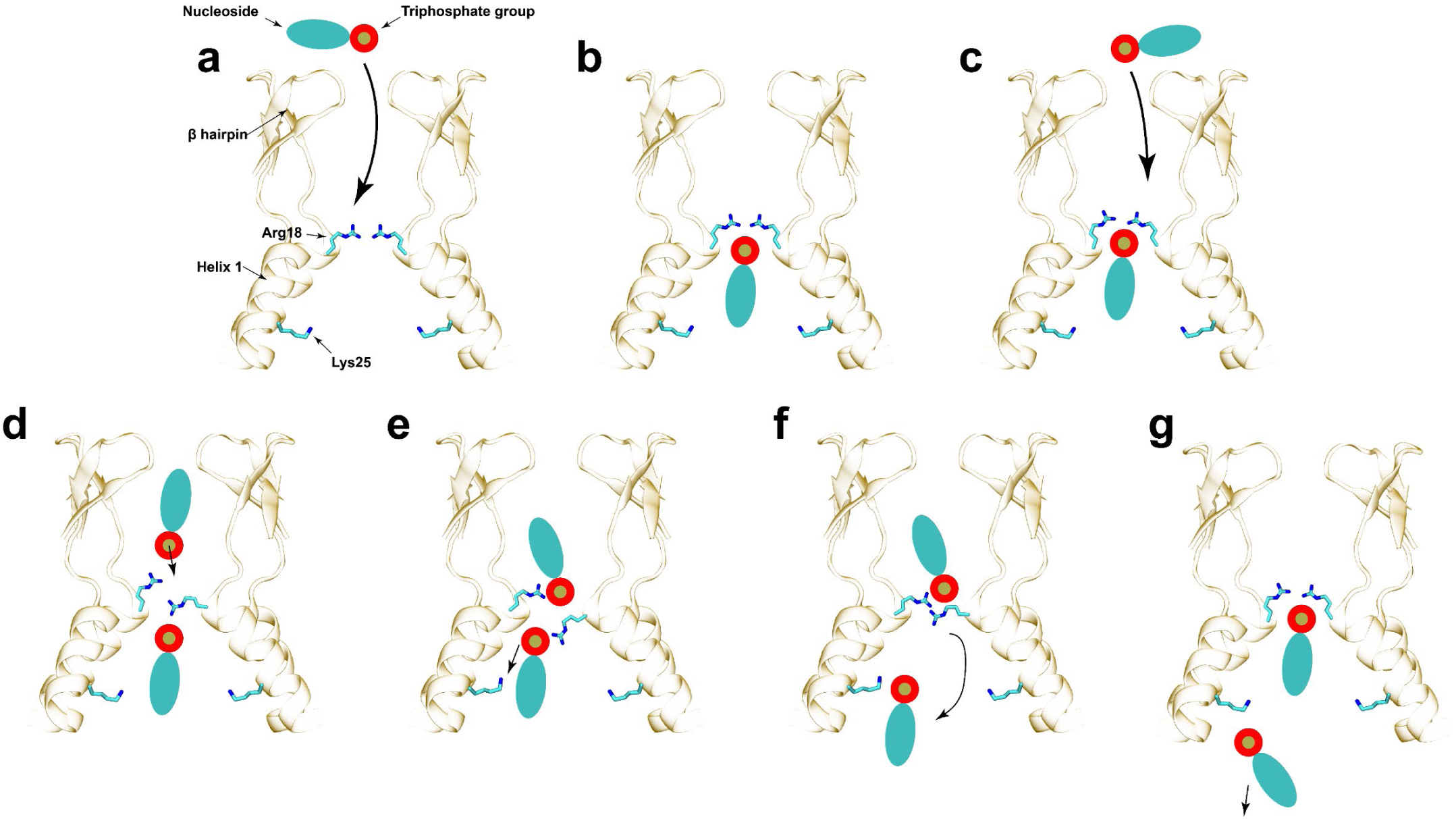
Molecular mechanism for nucleotide translocation through the HIV-1 CA hexamer. (a) A nucleotide diffuses freely between the exterior (top) of the capsid and the central cavity. (b) Subsequently, the nucleotide binds to Arg18 and Lys25 in a canonical binding conformation. Exact interactions between the nucleoside and helix-1 confer nucleotide-type specificity. (c) A second nucleotide freely diffuses between the exterior and the central cavity of CA. (d) The phosphate group of the second nucleotide interacts with Arg18 delocalizing the interactions between the first nucleotide and the Arg18 ring. (e) The second nucleotide enhances interactions between Lys25 and the first nucleotide. (f) Interactions between Lys25 and the phosphate group are weak and thermal fluctuations facilitate dissociation of the dNTP into the interior of the capsid. (g) The second nucleotide now occupies the canonical binding position (b) for a single nucleotide in the cavity.

Ion channels and transporters have evolved to employ a cooperative translocation mechanism to displace cargos between distinct environments^28,29^. Our free energy calculations for multiple small molecules suggest that binding of two nucleotides within the CA cavity promotes dNTP import. Interestingly, similar to water molecules in aquaporins^29^, dNTPs flip as they pass through hexamers. Furthermore, pure electrostatic interactions between CA and dNTPs are insufficient to describe the molecular mechanism of dNTP import, as the loss and gain of entropy associated with dewetting dNTPs and Arg18 is a key contributor to the free energy of binding. As a result, the cooperative mechanism for nucleotide translocation is not universal for any set of negatively charged molecules, as IP_6_ promotes dNTP import while BHC inhibits it.

We conclude that HIV-1 has evolved to import dNTPs from the cytoplasm into the capsid and to discern nucleoside type. Molecular determinants of this nucleoside-specific importing mechanism are encoded in highly conserved residues such as Lys25. In addition, we conclude that IP_6_ enhances import of dNTPs for efficient reverse transcription. The discovery of metabolite-dependent nucleoside-specific import provides a unique target for development of new therapies against HIV-1.

## Methods

### CA hexamer model building

The initial coordinates of the HIV-1 CA hexamer (Fig. S8a, b) were generated by applying a six-fold symmetry operation onto a native full-length HIV-1 CA (PDB accession code 4XFX). The two loops between residues 5 to 9 and residues 222 to 231, missing in the original structure, were built using Modeler^26^. Once the hexamer was built, the protonation states of titratable residues, namely histidine, asparagine, lysine and cysteine, were assigned using PDB2PQR^30^ at pH 7.4.

### Molecular mechanics parameterization of NTPs, IP_6_ and BHC

Parameters for the negatively-charged small molecules (Fig. S9), except ATP, which has parameters available in the CHARMM general force field, were derived by analogy following the CGENFF protocol^31^. The parameter penalties and charge penalties in each generated parameter files were less than ten indicating good analogy with the available atom types present in CGENFF^32^. A magnesium ion was added to the triphosphate group present in dNTPs. The coordination number of the magnesium ion with the phosphate group of the dNTP was constrained using *coordnum* in Colvars^33^.

### Simulation setup

Small charged molecules such as NTP/IP_6_/BHC were placed in the central cavity of the CA hexamer model. The models were then solvated using the TIP3P water model ^34^. Additionally, excess of TIP3P water molecules were deleted to transform the cubic water box into a hexagonal orthorhombic cell of dimension 92.328 Å in the 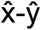 -ŷ plane and 90 Å in the 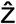 direction. The length of the system in the 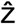 direction provided sufficient solvent padding, greater than 24 Å, to avoid interactions between periodic images. Na and Cl ions were then added to neutralize the system and the bulk salt concentration was set to the physiological concentration of 150 mM. The total number of atoms of the resulting CA hexamer models was 60,000 (Fig. S8c, d). Solvated WT and K25A, K25N, K25E and K25E/E28K CAs were derived using the mutator plugin in VMD.

### System minimization and equilibration

The solvated systems were then subjected to minimization in two stages, both using the conjugated gradient algorithm^35^ with linear searching^36^. Each stage consisted of 10,000 steps of energy minimization. During the first stage, only water molecules and ions were free to move, while the CA protein and NTP/IP_6_/BHC molecules were fixed. In the second stage, the backbone atoms of the CA protein were restrained with a force constant of 10.0 kcal mol^-1^ Å^-2^. Convergence of the minimization procedure was confirmed once the variance of the gradient was below 0.1 kcal mol^-1^ Å^-1^. Following minimization, the systems were tempered from 50 K to 310 K in increments of 20 K over 1 ns. Subsequently, the systems were equilibrated at 310 K for 100,000 steps, while the protein backbone atoms were restrained. Then positional restraints were gradually released at a rate of 1.0 Kcal mol^-1^ Å^-2^ per 400 ps from 10.0 Kcal mol^-1^ Å^-2^ to 0.0 Kcal mol^-1^ Å^-2^. All MD simulations were performed using NAMD 2.12^37^ with the CHARMM36m force field^38^. An internal time step of 2 fs was employed in the multi-step r-RESPA integrator as implemented in NAMD, bonded interactions were evaluated every 2 fs, and electrostatics were updated every 4 fs. Temperature was held constant at 310 K using a Langevin thermostat with a coupling constant of 0.1 ps^-1^. Pressure was controlled at 1 bar using a Nose-Hoover Langevin piston barostat with period and decay of 40 ps and 10 ps, respectively. The Shake algorithm was employed to constraint vibrations of all hydrogen atoms. Long range electrostatics was calculated using the particle-mesh-Ewald summation with a grid size of 1 Å and a cutoff for short-range electrostatics interactions of 12 Å.

### Gibbs free energy calculations

Progress variables (PVs), akin to reaction coordinates, for all free-energy molecular dynamics calculations were chosen as the location on the Z axis of the center of mass of the small-molecule(s) of interest. The origin of the progress variable was set to the center of mass of C_α_ atoms in the N-terminal domain of the CA hexamer (Fig. 1c). One- and two-dimensional potentials of mean force (PMFs) along the PVs were calculated using the Hamiltonian Replica-exchange/Umbrella Sampling (HREX/US) method^35^ implemented in NAMD 2.12^31,39^. The initial coordinates for the HREX/US windows were derived from constant-velocity steered MD (SMD) simulations in which molecules were pulled along the PV at a rate of 0.1 nm/ns. The center of mass of the small-molecules were positionally resta rained in the US windows with a harmonic force constant of 2.5 Kcal mol^-1^ Å^-2^ using Colvars^40^. The width of all US windows was set to 0.75 Å; except for single nucleotide HREX simulations (simulation #1-8) in which the window width was set to 1.0 Å. The number of US windows in the present HREX simulations were chosen so that the nucleotide translocation from capsid exterior to interior through the central cavity was uniformly sampled (Fig. 1c).

Two dimensional (2D) HREX/US simulations were employed to study the cooperativity between small molecules for translocation. For this purpose, two progress variables were employed: PV1 that determined the location of dATP, and PV2 that determined the location of dATP, IP_6_ or BHC in the CA cavity. The initial configurations for the 2D simulations were derived by pulling dATP using constant velocity SMD at 0.1 nm/ns along PV1, while the center of mass of dATP/IP_6_/BHC was restrained with a harmonic force constant of 2.5 Kcal mol^-1^ Å^-2^ using Colvars (cycle 1, Fig. S10a-c). A seeding method similar to that reported^41^ was employed to generate new simulation windows along PV2 as follows. For each conformation resulting from the SMD simulation, two new seeds were generated by displacing by ±0.75 Å the harmonic restraints for dATP/IP_6_/BHC along PV2 (cycle 2, Figure S10a-c). Subsequently, 5 ns of HREX/US MD simulation were performed. The last configuration from each replica was then used as the initial seed for a subsequent cycle where the harmonic restraints were displaced by another ±0.75 Å along PV2 (cycle 3, Figure S10a-c). The seeding process was repeated one last time and production 2D HREX/US simulations (production runs, Figure S10a-c) were performed for 30 ns per window. During each HREX/US simulation, exchange of the harmonic potential between neighboring replicas was attempted every 1000 steps (2 ps). Exchanges were attempted randomly between successive replicas; trial attempts for all replicas were equally probable. The following Metropolis Monte Carlo exchange criterion^42^ was employed

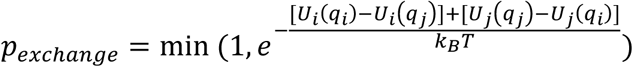

Where T=310 K, *k*_*B*_ is the Boltzmann constant, *q*_*i*_ and *q*_*j*_ denote the 3D conformations for two replicas i and j, and *U*_*i*_ and *U*_*j*_ represent the potential energies derived from the restrained Hamiltonian evaluated at the indicated conformation.

Potentials of mean force were derived from the resulting sampling in each of the US windows using the weighed histogram analysis method (WHAM)^43^. In WHAM, PMF bins are obtained from US/HREX simulation windows. Convergence of the 1D US/HREX calculations was characterized by the changes in the resulting PMF in trajectory increments of 10 ns (Fig. S11). That is, simulations were considered to have converged once the maximum change in one of the PMF bins resulting from adding more simulation data, was less than 1 Kcal mol^-1^.

Convergence of the 2D calculations was examined by means of the fluctuations of the average root mean squared error for each PMF bin (RMSE) (Fig. S12a). Each 2D HREX/US simulation was divided into 60 time-slices of width 0.5 ns. Then the PMF was calculated for each slice using WHAM. Using the last slice as the reference, the RMSE of the *k-*th slice was calculated using the equation:

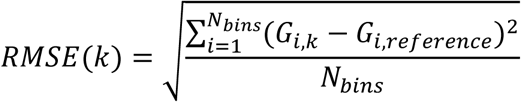

Where the sum runs over the total number of PMF bins (*N*_*bins*_), G_*i, k*_ and G_*i, reference*_ are the free energies at the *i*-th PMF bin for the *k*-th and last slice, respectively. Convergence of 2D PMF was established once the RMSE was not larger than 3.0 kcal/mol. The bins before convergence were discarded. Subsequently, utilizing the converged slices, the 2D PMF surfaces (Fig. 2) and their standard deviations (Fig. S12b, c and d) were computed using WHAM. The statistics of the exchange ratios between neighboring US replicas in HREX simulations are shown in Fig. S11. Overall, the exchange ratios are greater than 20% in 2D HREX simulations, indicating enough sampling efficiencies (Fig. S13).

### Cell lines

HEK 293T, GHOST and HeLa cells were maintained at 37°C in Dulbecco’s modified Eagle medium supplemented with 10% heat-inactivated bovine serum, 1% penicillin-streptomycin, and 1% glutamine in 5% CO_2_. Human PBMCs were isolated from leukopheresis packs obtained from the Central Blood Bank (Pittsburgh, PA) via Ficoll-Paque Plus (GE Healthcare) density gradient centrifugation, following manufacturer’s instructions, and stimulated with 50 U/mL recombinant interleukin-2 (IL-2, Thermo Fisher) and 5 μg/mL phytohaemagglutinin (PHA, Gibco) for 48-72h prior to infection. CD14+ monocytes were purified from PBMCs via CD14 MicroBeads (Miltenyi) magnetic cell sorting, following manufacturer’s instructions, and differentiated into macrophages with 50 ng/mL granulocyte macrophage colony-stimulating factor (GM-CSF, R&D Systems) for 6d prior to infection.

### Isolation of HIV-1 capsids for AFM measurements

HIV-1 pseudovirions were used to isolate capsids as previously described^44,45^. Briefly, approximately 10^6^ human embryonic kidney (HEK) 293T cells were transfected with 2.5 μg of ΔEnv IN-HIV-1 plasmid (DHIV3-GFP-D116G)^11^ using 10 μg of polyethylenimine (PEI, branched, MW ∼25,000, Sigma-Aldrich). After 20 h, the medium was replaced with fresh medium. After 6 h, the supernatant was harvested, centrifuged at 1,000 rpm for 10 min, and filtered through a 0.45-μm-pore size filter. The virus-containing supernatant was concentrated by ultracentrifugation in a SW-28 rotor (25,000 rpm, 2 h, 4°C) using OptiPrep density gradient medium (Sigma-Aldrich). Virus containing fractions were collected, mixed with 10 ml of TNE buffer (50 mM Tris-HCl, 100 mM NaCl, 0.1 mM EDTA, pH 7.4) and added to 100-kDa molecular mass cutoff Vivaspin 20 centrifugal concentrators (100,000 MWCO, Sartorius AG, Germany). The mixture was centrifuged twice at 2,500 × g for 25 to 30 min at 4°C, until the supernatant level in the concentrators reached 200–300 μL.

Capsids were isolated from concentrated virus-containing material using a previously described protocol^46^ with modifications. Briefly, ∼40 μL of purified HIV-1 pseudovirions was mixed with an equal amount of 1% Triton-X diluted in 100 mM 3-(N-morpholino) propane sulfonic acid MOPS buffer (pH 7.0) and incubated for 2 min at 4 °C. The mixture was centrifuged at 13,800 × g for 8 min at 4°C. After removing the supernatant, the pellet was washed twice by adding ∼80 μl of MOPS buffer and centrifuging at 13,800 × g for 8 min at 4°C. The pellet was resuspended in 10 μl of MOPS buffer.

### AFM measurements and analysis

AFM measurements and analysis were performed as previously described^11,24^. Briefly, 10 μL of isolated HIV-1 capsids resuspended in MOPS buffer was incubated for 30 min at room temperature on hexamethyldisilazane (HMDS)-coated microscope glass slides. Measurements were carried out in MOPS buffer, without sample fixation. Every experiment was repeated at least three times, each time with independently purified HIV-1 capsids. IP_6_ was provided from the James laboratory as a generous gift. Measurements were carried out with a JPK Nanowizard Ultra-Speed atomic force microscope (JPK Instruments, Berlin, Germany) mounted on an inverted optical microscope (Axio Observer; Carl Zeiss, Heidelberg, Germany) using silicon nitride probes (mean cantilever spring constant, kcant=0.12 N/m, DNP, Bruker). Height topographic and mechanical map images were acquired in quantitative imaging (QI) mode, at a rate of 0.5 lines/s and a loading force of 300 pN. All measurements were carried out in MOPS buffer which contained 100 μM IP_6_ or mellitic acid (Sigma) when relevant.

Capsid stiffness was obtained by the nanoindentation method as previously described^11,24,46^. Briefly, the stiffness value of each capsid was determined by acquiring ∼400 force-distance (F-D) curves. To determine the stiffness value of capsids, 20 F-D curves at rate of 20 Hz at each of 24 different points on the capsid surface were determined. To confirm that the capsid remained stable during the entire indentation experiment, we monitored individual measured point stiffness as a histogram (Fig. S14a) and as a function of the measurement count (Fig. S14b). Samples whose point stiffness values decreased consistently during experimentation were discarded, since they underwent irreversible deformation.

The maximum indentation of the sample was 4 nm, which corresponds to a maximum loading force of 0.2 to 1.5 nN. Prior to analysis, each curve was shifted to set the deflection in the noncontact section to zero. The set of force distance curves was then averaged (Fig. S14c). From the slope of the averaged F-D curve, measured stiffness was derived mathematically. The stiffness of the capsid was computed using Hooke’s law on the assumption that the experimental system may be modeled as two springs (the capsid and the cantilever) arranged in series. The spring constant of the cantilever was determined during experiment by measuring thermal fluctuation^47^. To reduce the error in the calculated point stiffness, we chose cantilevers such that the measured point stiffness was <70% of the cantilever spring constant. Data analysis was carried out using MATLAB software (The Math Works, Natick, MA).

### Viruses

Mutations were generated in previously-described single-round plasmid derivatives of HIV-1_NL4-3_ (pNLdE-luc)^48^ and HIV-1_LAI_ (pBru3ori-ΔEnv-luc2)^49^ that encoded for luciferase via the QuikChange site-directed PCR mutagenesis kit (Agilent). Resulting plasmid DNAs were verified by Sanger sequencing. Virus was produced in HEK 293T cells via transfection of the proviral plasmids with pL-VSV-G^50^ and, for imaging assays, pcDNA5-TO-Vpr-mRuby3-IN^51^ using Lipofectamine 2000 (Invitrogen). Supernatants were harvested after 48 h, centrifuged to remove cells, and stored in aliquots at −80° C. The infectivity of HIV-1 stocks normalized by p24 content (XpressBio) was assayed by infection of GHOST cells as previously described^52^.

### ERT with WT and CA-mutant protein for AFM analysis

Reverse transcription was induced in isolated capsids attached to HMDS-coated microscope glass slides. To initiate reverse transcription, MOPS buffer was replaced with reverse transcription buffer (100 µM dNTPs and 1 mM MgCl_2_ in 100 mM MOPS buffer, pH 7.0) ^25^. All measurements were carried out at room temperature (23-25°C).

### HIV-1 CA retention assay

Equal p24 amounts of virus containing mRuby3-IN were diluted in STE buffer (100 mM NaCl, 10 mM Tris-HCl, 1 mM EDTA, pH 7.4) supplemented with 100 μM IP_6_, adhered to 35 mm glass-bottom dishes (MatTek), coated with Cell-Tak (Corning) for 30 min at 37°C following manufacturer’s instructions, washed with PBS and lightly permeabilized with 0.02% Triton X-100 (Fisher Scientific) for 30 sec prior to washing and fixation with 2% paraformaldehyde (PFA). For IP_6_ studies, samples that had been adhered to glass and lightly permeabilized as above were additionally incubated for 2 h at 37°C in STE buffer with or without 100 μM IP_6_, and lightly permeabilized again after incubation prior to washing and fixation. Samples were permeabilized with 0.1% Triton X-100, blocked with normal donkey serum, immunostained for CA/p24 using mouse anti-CA monoclonal antibody 24-4 (Santa Cruz) and donkey anti-mouse IgG-Cy5, and mounted with coverslips. Confocal imaging was performed on a Nikon A1 laser scanning confocal microscope, with 6 fields imaged per sample. CA staining and IN particles were enumerated with Imaris imaging analysis software (Bitplane).

### Infectivity assays and measurement of reverse transcripts and 2-LTR circles

Cells were plated overnight in 24-well or 96-well plates and challenged with virus at equal amounts of p24. Virus infectivity was determined by luciferase production (Promega) after 48h using a Synergy2 Multi-Detection Microplate Reader (BioTek)

HeLa cells were plated in 6-well plates and infected with virus at equal amounts of p24 after treatment with DNaseI (Roche) for 30 min at 37°C. Control infections were performed in the presence of 25 nM rilpivirine. After 24 h, cells were washed with PBS, trypsinized, and pelleted, and DNA was extracted using the Blood Mini Kit (Qiagen). Early (RU5) and late (gag) HIV-1 reverse transcripts and 2-LTR circles were measured by qPCR as previously described^18^. Values obtained in the presence of rilpivirine were subtracted from experimental values to determine levels of viral reverse transcription and 2-LTR circle formation.

### Statistics of infectivity assays, quantitative PCR, and capsid stability assay

Results were analyzed for statistical significance by two-sided student t test (infectivity assays and quantitative PCR) or one-way ANOVA with Tukey’s multiple comparisons test (capsid stability assay) with Prism software (GraphPad). A p-value of less than or equal to 0.05 was used to indicate statistical significance.

### TEM of viruses

Virus produced from HEK 293T cells was centrifuged to remove cells, filtered through a 0.45 μm Polyethersulfone (PES) syringe filter (Millipore), transferred into 25×89mm polyallomer ultracentrifuge tubes (Beckman) and ultracentrifuged in an Optima XL-100K ultracentrifuge (Beckman Coulter) using an SW28 rotor at 25,500 rpm (117,250 x g) for 2.5 h at 4°C. Following aspiration of supernatant the pellet was fixed in FGP fixative (1.25% formaldehyde, 2.5% glutaraldehyde, and 0.03% picric acid in 0.1 M sodium cacodylate buffer, pH 7.4) for 2 h at room temperature and stored at 4°C. Ultrathin sections (60 nm) were cut on a Reichert Ultracut-S microtome, transferred to copper grids stained with lead citrate, and observed using a JEOL 1200EX microscope with an AMT 2k charge-coupled-device camera. Images captured at 30,000X magnification were visually inspected to classify viral particles as mature, immature, eccentric, or containing 2 apparent capsids, and over 100 particles were counted per virus preparation. Linear regression analysis of the eccentric particles was performed using GraphPad Prism software.

### Preparation of recombinant CA assemblies

U-^13^C,^15^N labeled CA K25N and K25A mutants were isolated and purified using the protocol reported previously with minor modifications^18^. Pure proteins were assembled as reported previously^11^ with minor modifications. First, the proteins were dialyzed overnight against 50 mM MES buffer pH 6.0 containing 300 mM NaCl. The dialyzed proteins were concentrated to 40 mg/mL and diluted to 1:1 volumetric ratio with assembly buffer (50 mM MES pH 6.0, 600 μM NaCl). The final assembly was carried at room temperature by 1:1 volumetric dilution with 1800 μM inositol hexaphosphate (IP_6_), pH 6.0. CA assemblies were pelleted after the incubation for 1 hour, and stored at 4 °C.

### Preparation of recombinant CA-SP1-NC assemblies

Protein was purified and assembled as previously described by the authors^46^.

### Transmission electron microscopy of CA assemblies

After assembly, a small aliquot of each sample was removed and immediately stained with 2% uranyl acetate on copper grids. Images were collected using the TALOS F200C at the Keck Center for Advanced Microscopy and Microanalysis of Interdisciplinary Science and Engineering (ISE) Laboratory of the University of Delaware.

### MAS NMR Spectroscopy of CA assemblies

MAS NMR experiments were performed on a Bruker 20.0 T narrow bore AVIII spectrometer. The Larmor frequencies of ^1^H, ^13^C, and ^15^N were 850.4, 213.8 and 86.2 MHz, respectively.

The U-^13^C, ^15^N-CA WT and K25A conical assemblies were packed into 3.2 mm thin wall MAS NMR rotors, and ^13^C-^13^C CORD data sets were acquired using a 3.2 mm E-Free HCN probe. The sample temperature was 5±1 °C controlled by the Bruker VT controller. The MAS frequency was 14.000±0.001 kHz controlled by the Bruker MAS-3 controller. The 90° pulse lengths were 2.9 μs (^1^H) and 4.3 μs (^13^C). The cross-polarization contact time was 1 ms; a linear 90-110% amplitude ramp of was applied on ^1^H; the center of the ramp was at 82 kHz and Hartmann-Hahn matched to the first spinning sideband; ω_RF_=54 kHz was applied on the ^13^C channel. SPINAL-64 ^1^H decoupling^53^ (ω_RF_=86 kHz) was applied during t_1_ and t_2_ periods. The CORD mixing time was 50 ms. The spectra were acquired as a 2048×864 (t_2_×t_1_) points complex matrix to the final acquisition times of 15.98 and 10.29 ms, respectively. States-TPPI protocol^54^ was used for frequency discrimination in the indirect dimension. A total of 156 transients were added up for each FID; the recycle delay was 2 s.

The U-^13^C, ^15^N-CA K25N conical assemblies were packed into a 1.3 mm MAS NMR rotor, and ^13^C-^13^C RFDR spectrum was acquired using a 1.3 mm HCN probe. The sample temperature was 5±1 °C controlled by the Bruker VT controller. The MAS frequency was 60.000±0.001 kHz controlled by the Bruker MAS-3 controller. The 90° pulse lengths were 1.4 μs (^1^H) and 2.85 μs (^13^C), with T_CP_ of 1 ms; a linear 90-110% amplitude ramp of was applied on ^1^H; the center of the ramp was at 100 kHz and Hartmann-Hahn matched to the first spinning sideband; ω_RF_=20 kHz was applied on the ^13^C channel. TPPM ^1^H decoupling^55^ (ω_RF_=10 kHz) was applied during t_1_ and t_2_ periods. The spectrum was acquired as a 2048×800 (t_2_×t_1_) points complex matrix to the final acquisition times of 15.98 and 9.84 ms, respectively. States-TPPI was used for frequency discrimination in the indirect dimension. A total of 256 transients were added up for each FID; the recycle delay was 2 s.

The spectra were processed with 50° shifted squared sine-bells in both dimensions and zero-filled to at least twice the number of points in the indirect dimension. The processed spectra were analyzed in NMRFAM-Sparky^56^. Chemical shifts of CA NL_4-3_ tubular assemblies^57^ were used for data analysis.

## Acknowledgements

The authors thank Angela Gronenborn for enlightening discussions and Teresa Brosenitsch for editorial support. This work was supported by the US National Institutes of Health (NIH) (P50AI1504817, R01AI147890, R01AI070042, R01GM107013) and by the Israel Science Foundation (Grant 234/17). We acknowledge NIH grant P30GM110758 for supporting the core instrumentation infrastructure at the University of Delaware. This work used the Extreme Science and Engineering Discovery Environment, which is supported by the National Science Foundation (Grant ACI-1548562). This work used XSEDE Bridges and Stampede2 at the Pittsburgh Super Computing Center and Texas Advanced Computing Center, respectively, through allocation MCB170096. This research also used resources from the Oak Ridge Leadership Computing Facility (OLCF) at Oak Ridge National Laboratory, which is supported by the Office of Science of the Department of Energy under Contract DE-AC05-00OR22725. This work also utilized microscopes in the University of Pittsburgh Center for Biologic Imaging (CBI). We acknowledge a Director’s Discretionary award on the Summit supercomputer from the OLCF.

## Author Contributions

J.R.P. designed and C.X. performed all-atom MD simulations. D.K.F. designed and performed all virus experiments. I.R. designed and S.R. performed capsid isolation and AFM experiments. J.A., B.R. and R.A.D. performed protein purification and/or in vitro assembly. B.R., W.L, A.N.E. and R.A.D. performed TEM and analyzed associated data. R. Z. and T.P. designed and performed NMR experiments. The manuscript was written primarily by J.R.P., D.K.F., C.A., C.X., I.R., T.P., A.N.E. and Z.A, with input from all authors. The project was originally conceived by J.R.P., with input from all authors throughout experimentation and manuscript preparation.

---

**Figure S1.**
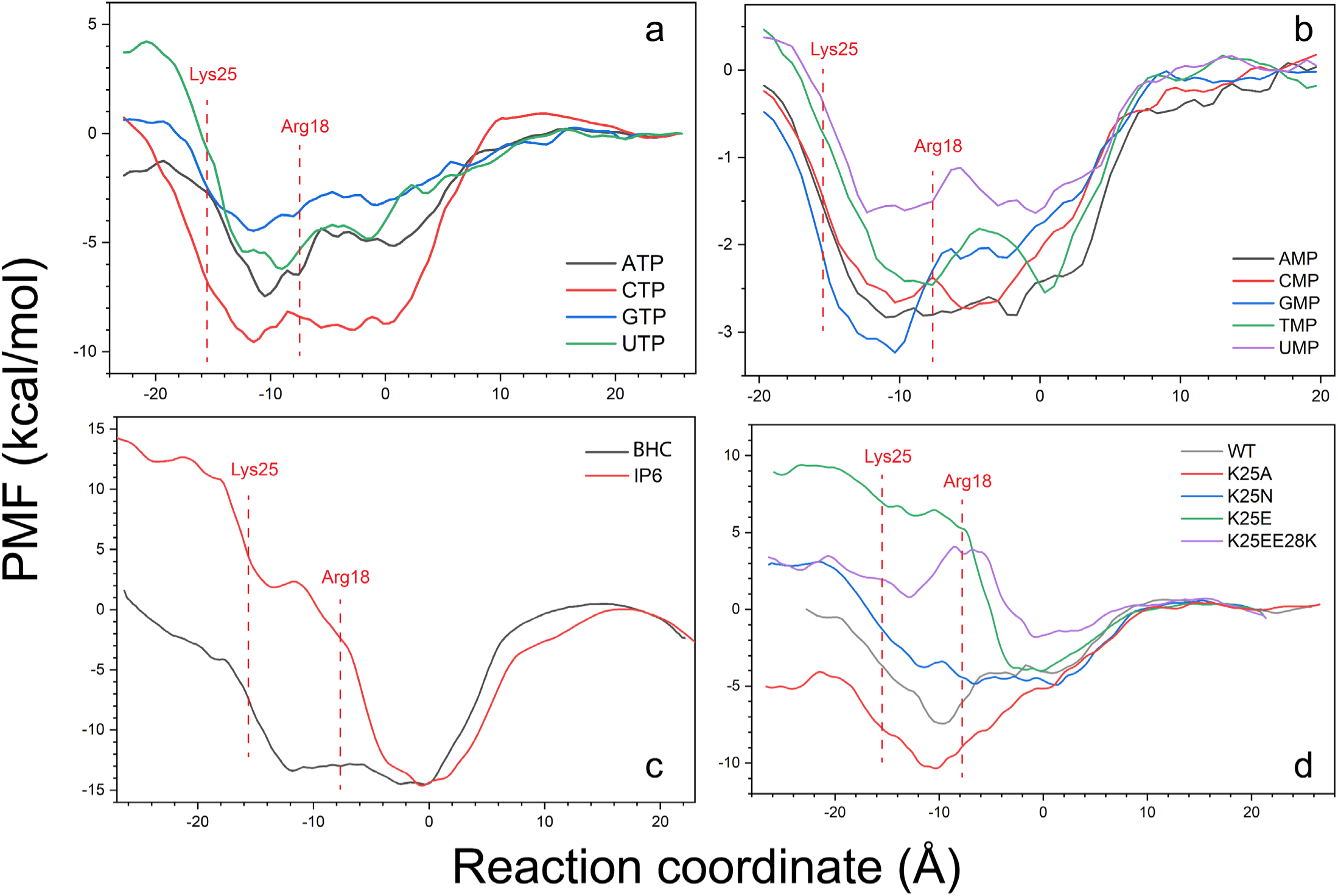
The 1D free energy landscapes of small molecules interacting with CA hexamers. (a) Binding profiles of rNTPs show different binding affinities and specifics compared to dNTPs. (b) Binding profiles of nucleotide monophosphates, which are weaker compared to rNTPs and dNTPs. (c) Binding profile of BHC and IP_6_ to a hexamer. BHC presents a much wider well compared to IP_6_. (d) 1D free energy landscapes of dATP binding to WT CA and indicated mutants. Compared to WT, K25A should result in leaky capsids, in contrast to K25N, K25E and K25E/E28K, which each should result in blocked capsid cavities.

**Figure S2.**
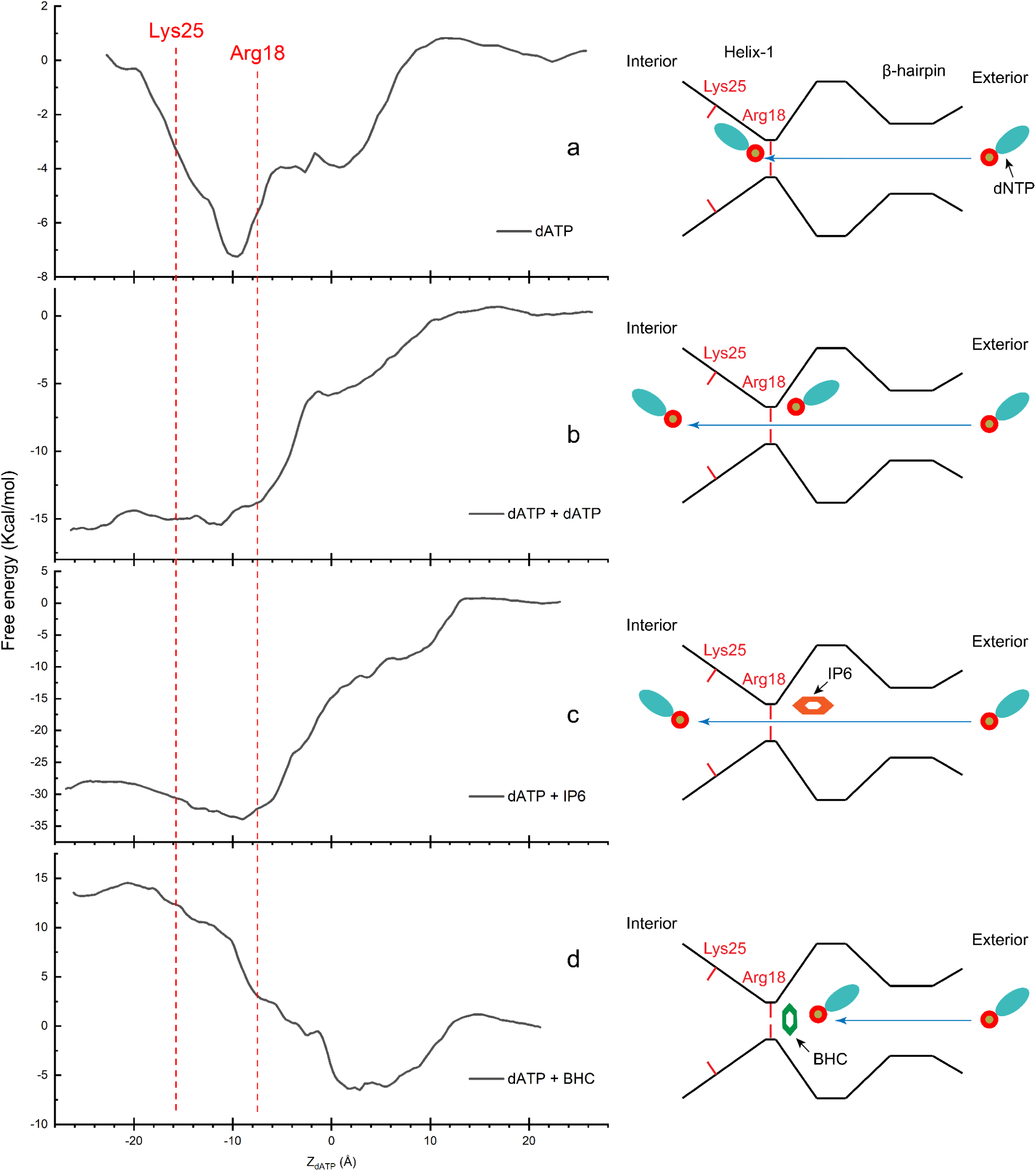
Free energy landscape of cooperative interactions between small molecules and the CA hexamer. (a) Free energy profile for a single dATP interacting with CA. The profile indicates dNTP binding, but not translocation as the energy barrier is too high to be overcome by simple thermal fluctuations. (b) Cooperation between multiple dATP molecules shifts the free energy profile from binding to a gradient pointing towards the interior of the capsid. (c) Cooperation between IP_6_ and dNTPs also creates a gradient toward the interior, but almost twice as strong compared to dNTPs alone. (d) BHC inside of the CA cavity facilitates dNTP binding but not translocation.

**Figure S3.**
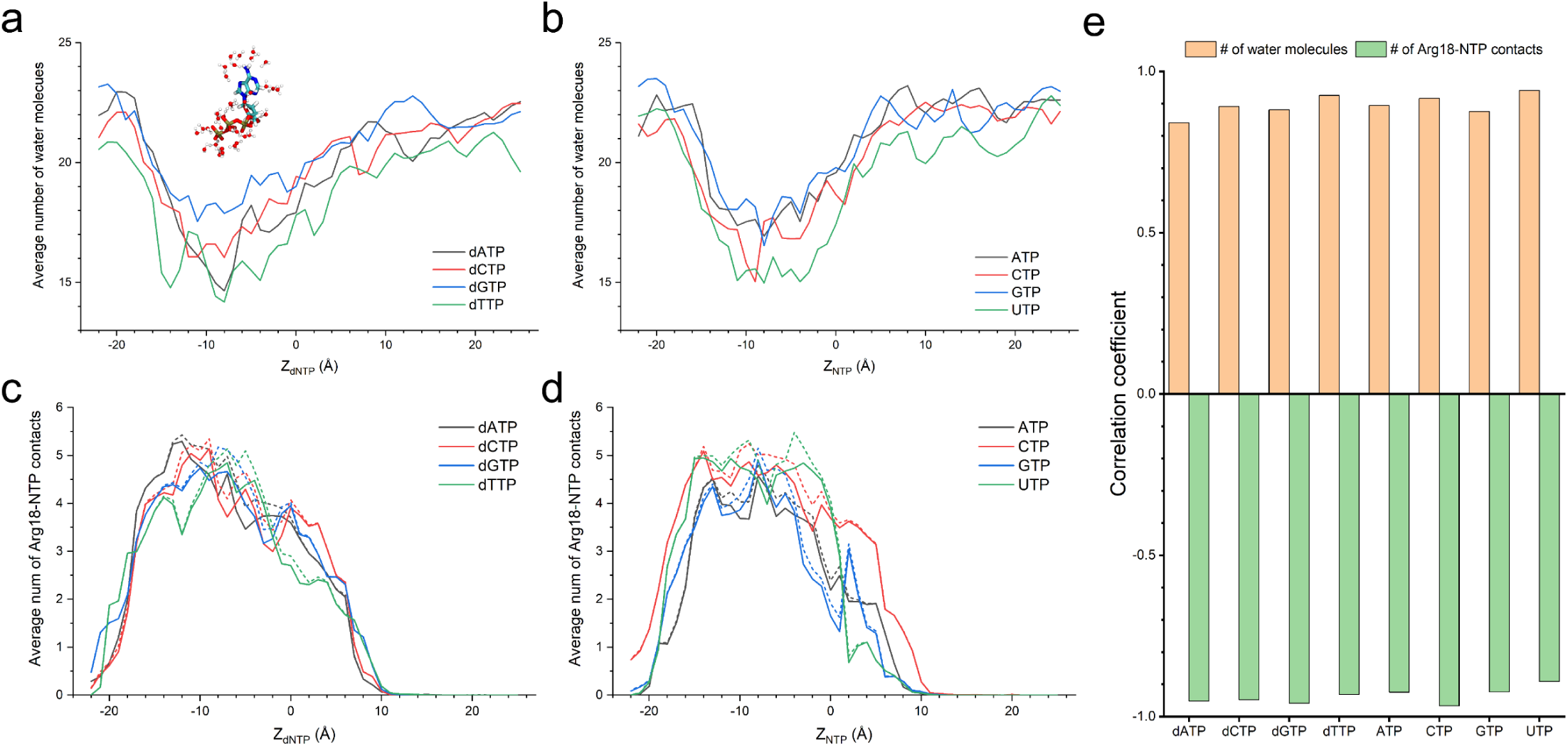
Mean number of water molecules interacting with dNTPs. (a) A significant de-wetting of dNTPs is observed near Arg18, indicating a loss of conformational entropy. (b) Similarly, loss of water molecules is observed for rNTPs. The loss of entropy is compensated by the formation of salt-bridges between Arg18 and dNTPs (c) and rNTPs (d). (e) Correlation analysis between loss of solvation molecules and formation of bonds with Arg18 and NTPs. Importantly, solvation of the dNTP/rNTP as it moves toward the interior of the capsid is assisted by several charged or polar residues including K25/30 and E28/29.

**Figure S4.**
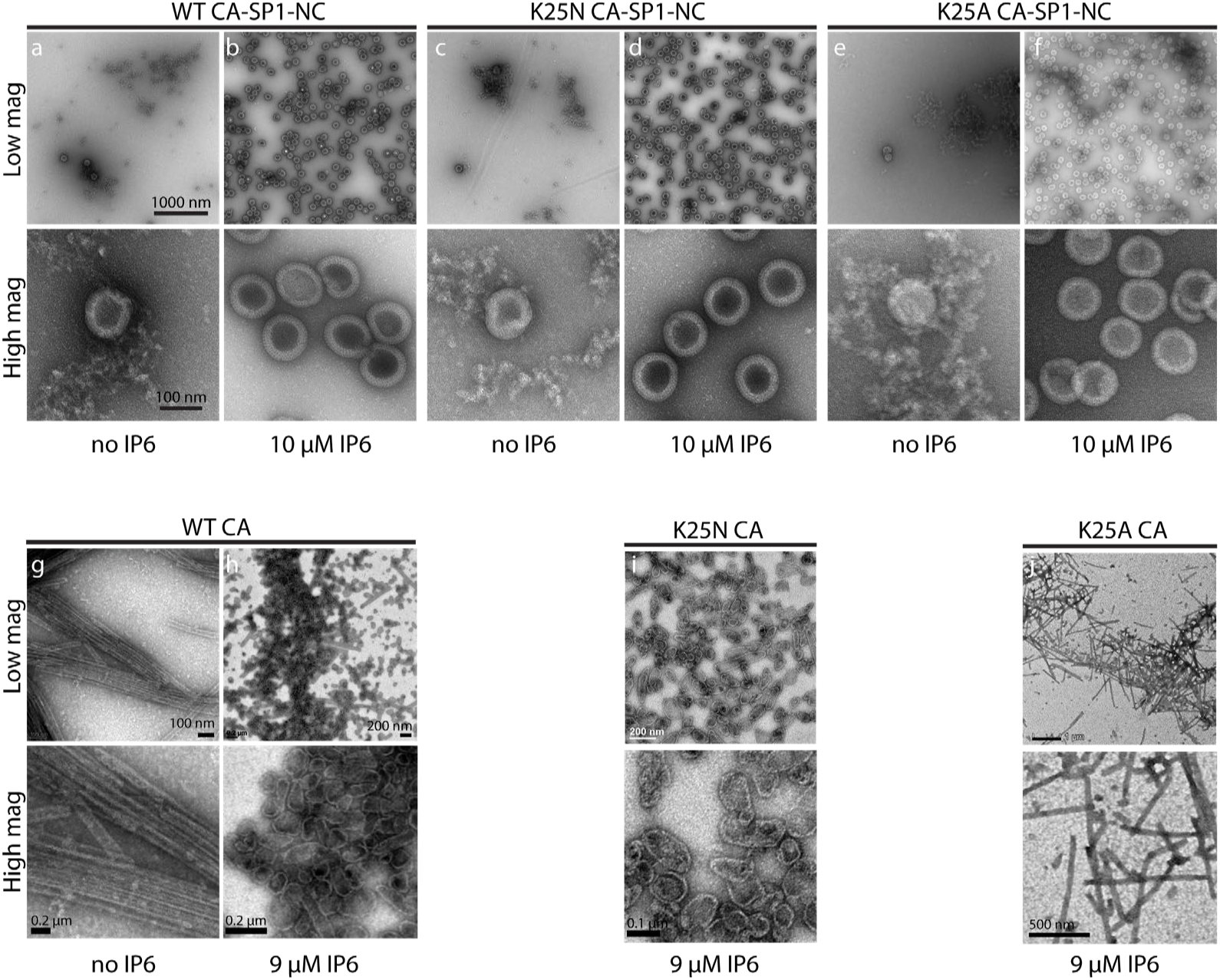
In vitro assembly of CA-SP1-NC and CA, NL4-3 strains. (a-f) Negatively stained TEM images of WT (a, b), K25N (c, d) and K25A (e, f) CA-SP1-NC. The mutants produce virus like particles with immature morphologies. Assemblies in (a, c, e) contain 2 mg/ml of the respective protein in 50 mM Tris-HCl (pH 8.0), 100 mM NaCl. Assemblies in (b, d, f) contain 10 mg/ml of the respective protein in 50 mM MES (pH 6.0), 10 μM IP_6_. (g-j) Negatively stained TEM images of WT (g, h) K25N (i) and K25N (j) CA show virus-like particles of mature morphology, namely tubes and cones. Assemblies in (g) contain 20 mg/ml CA in 50 mM Tris-HCl (pH 8.0), 2.4 M NaCl. Assemblies in (h, I, j) contain 20 mg/ml of the respective protein in 50 mM MES (pH 6.0), 0.9 μM IP_6_.

**Figure S5.**
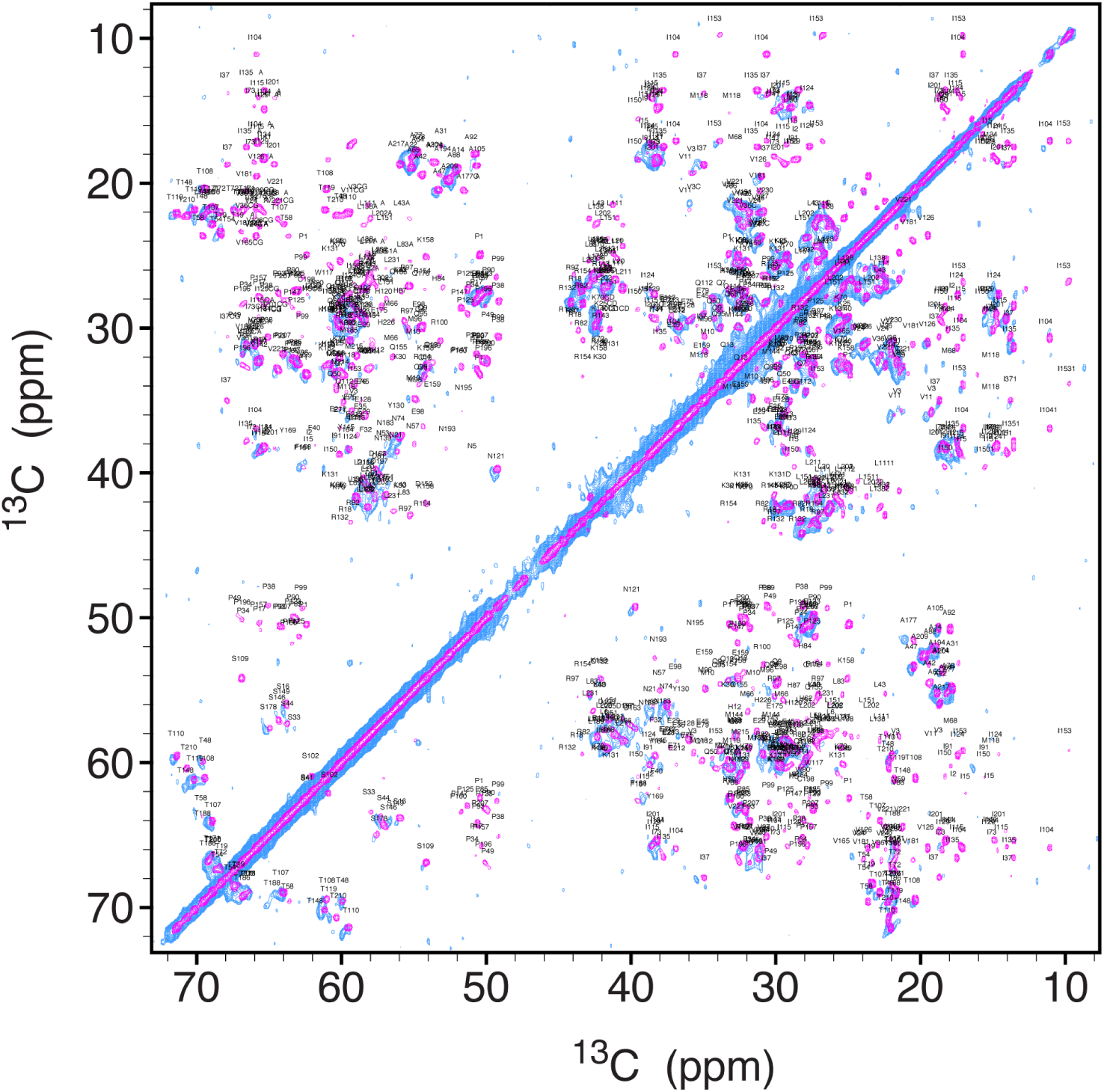
2D ^13^C-^13^C correlation MAS NMR spectra of conical assemblies of CA (CORD, magenta) and CA K25N mutant (RFDR, light blue). The assemblies contain 20 mg/ml of the respective protein in 50 mM MES (pH 6.0), 0.9 μM IP_6_. The similar chemical shifts indicate that K25N mutant is folded and its overall structure is the same as in the WT CA conical assemblies.

**Figure S6.**
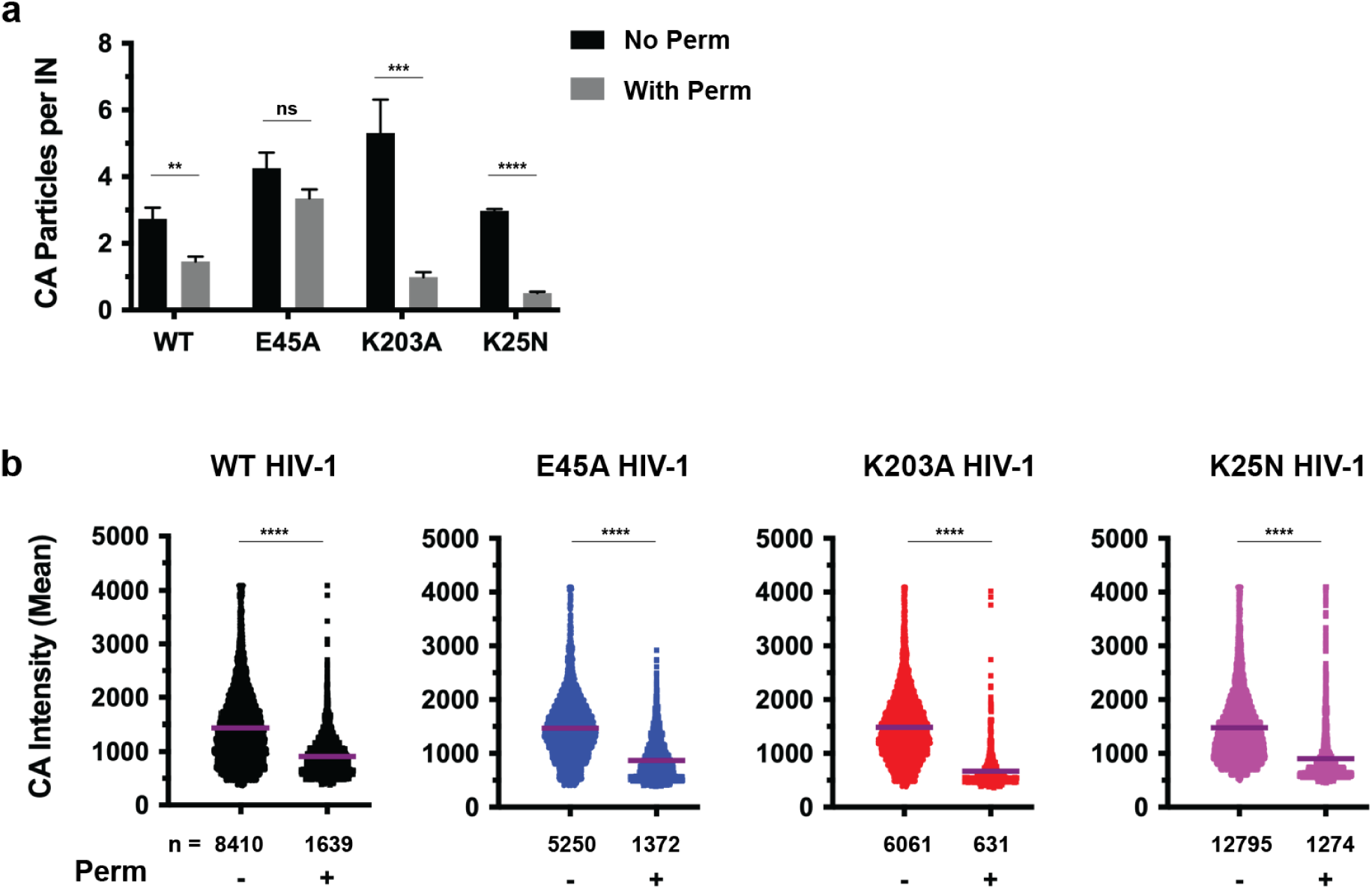
CA retention imaging assay distinguishes WT, hyperstable, and hypostable HIV-1 capsid phenotypes. WT or mutant viruses containing mRuby3-IN were captured on glass either with or without pre-fixation membrane permeabilization, immunostained for CA, and imaged. (A) The number of CA particles per IN particles is shown for each virus +/- permeabilization. E45A and K203A HIV-1 are included as examples of hyperstable and hypostable CA mutants, respectively^22^. Error bars indicate SEM for 2 experiments. (B) The mean fluorescence intensity of CA staining for each imaged virus particle is shown, with the overall mean intensity for the population indicated by the purple bar. Representative results are shown from one experiment. ** P < 0.01; *** P < 0.001; **** P < 0.0001.

**Figure S7.**
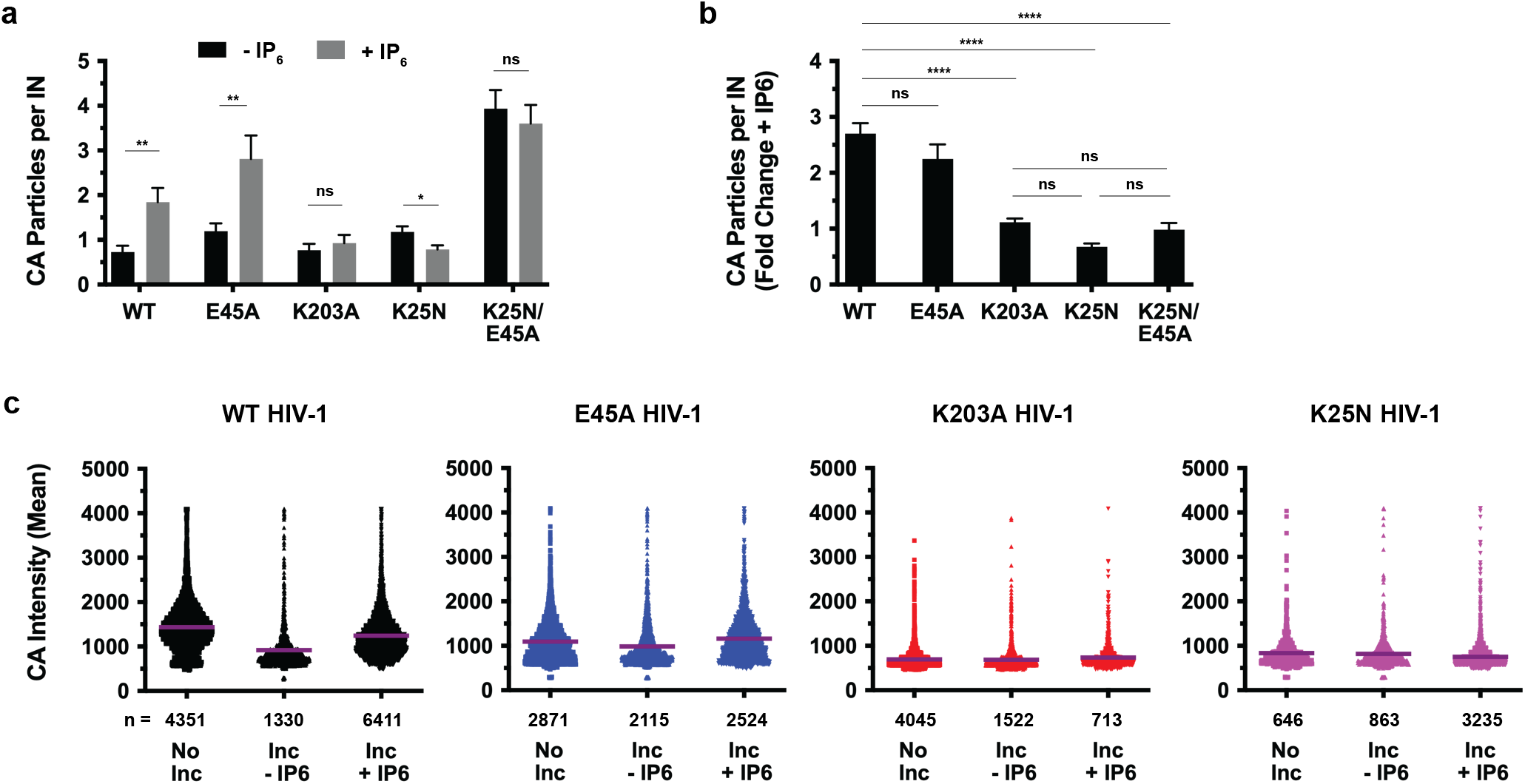
The effect of IP_6_ on HIV-1 CA retention. WT or mutant viruses containing mRuby3-IN were captured on glass, lightly permeabilized, and incubated for 2 h in STE buffer with or without 100 μM IP_6_. Viruses were lightly permeabilized again, fixed, immunostained for CA, and imaged. (A) The number of CA particles per IN particles is shown for each virus +/- IP_6_. Error bars indicate SEM for 2 experiments. (B) The data from (A) are shown expressed as the fold change of CA staining retained for each virus with the addition of IP_6_ during incubation. (C) The mean fluorescence intensity of CA staining for each imaged virus particle is shown, with the overall mean intensity for the population indicated by the purple bar. Representative results are shown from one experiment. * P < 0.05; ** P < 0.01; *** P < 0.001; **** P < 0.0001.

**Figure S8.**
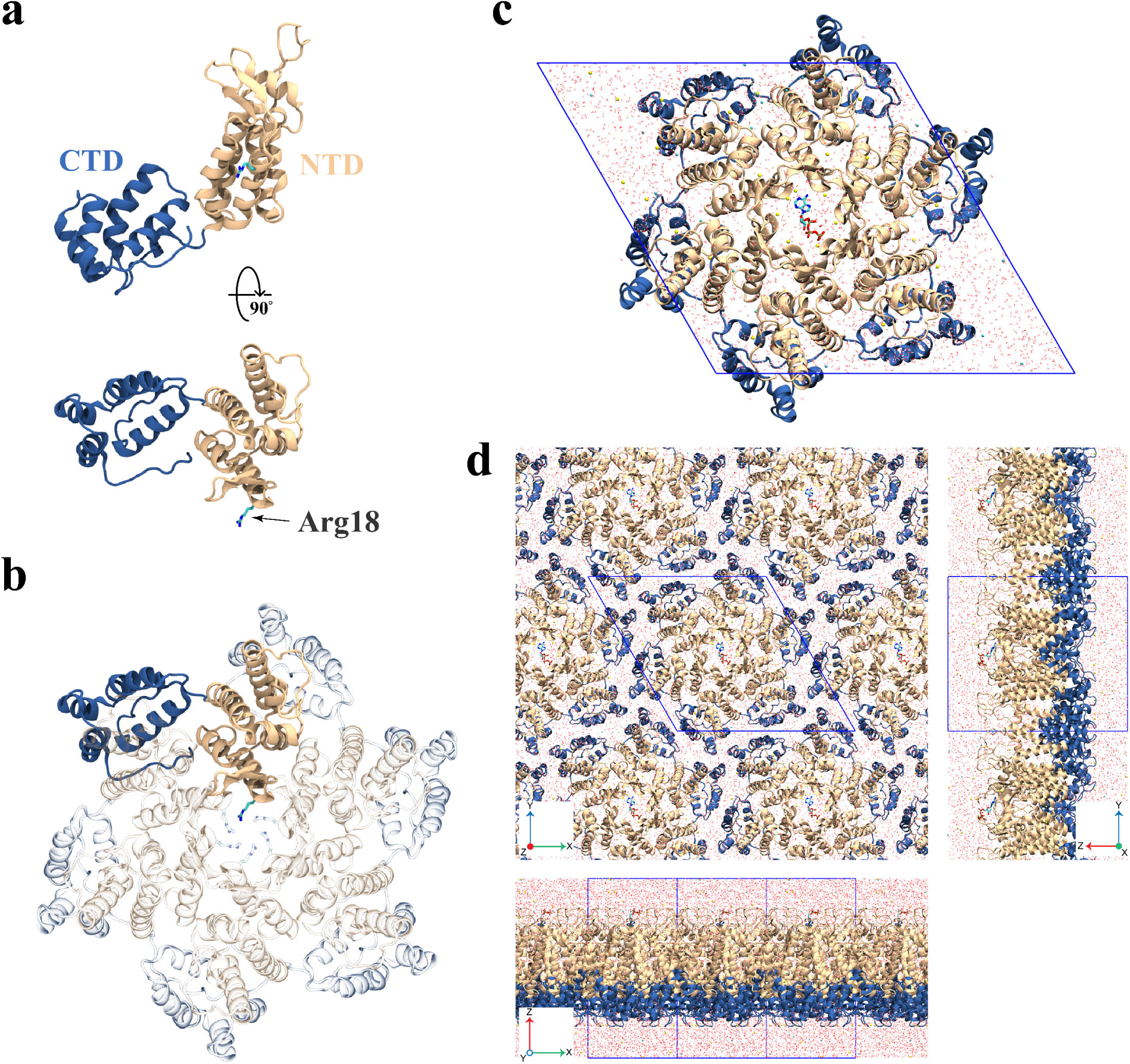
Atomistic models employed in the present study. (a) The structure of a HIV-1 CA monomer and (b) hexamer. (c) The CA hexamer in a hexagonal water box with 150 mM NaCl and dATP. (d) A flat CA hexamer lattice formed by applying periodic boundary conditions.

**Figure S9.**
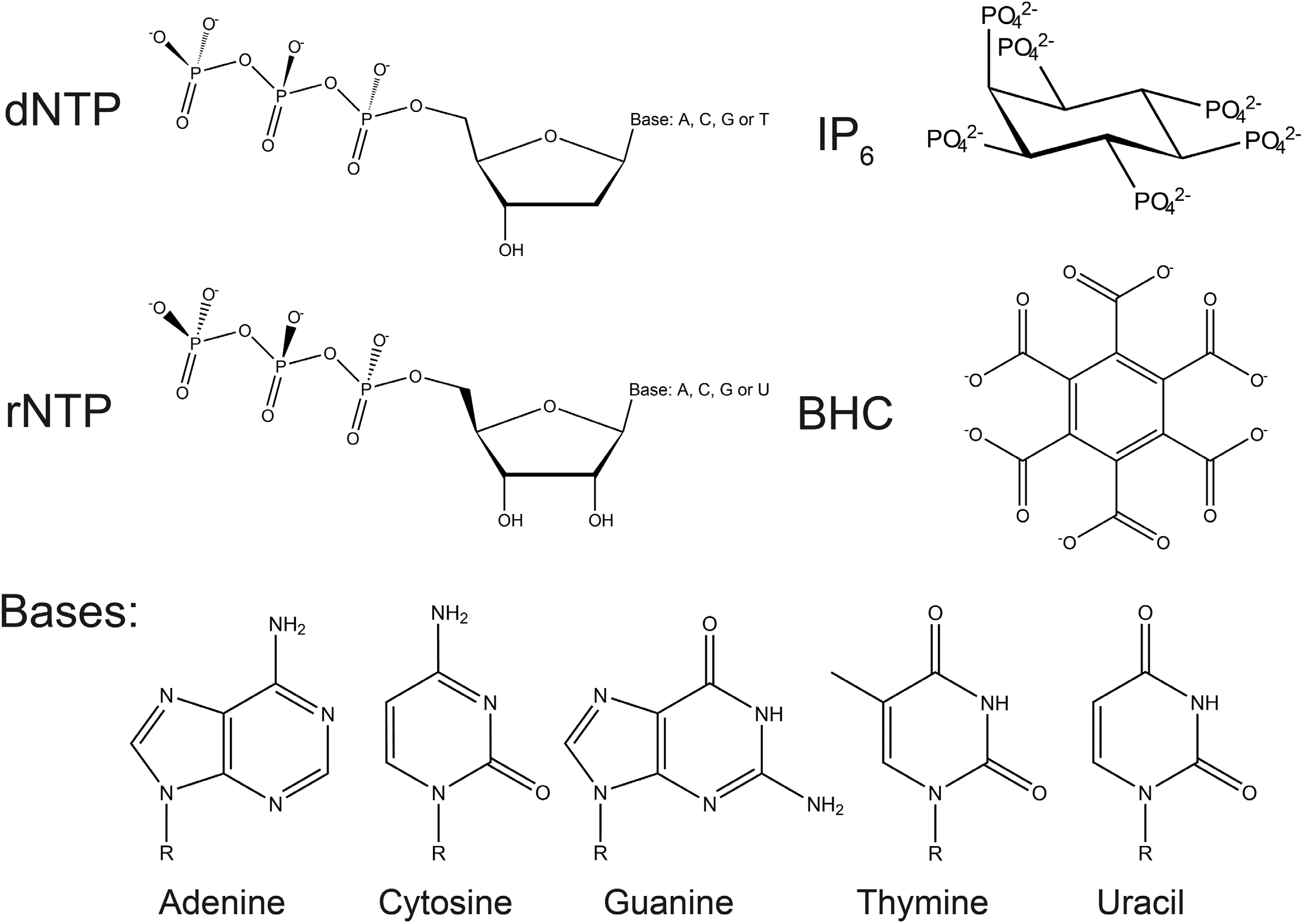
Molecular structures of small molecules used in the present study. dNTP, rNTP, myo-IP_6_, BHC, and dNTP and rNTP bases.

**Figure S10.**
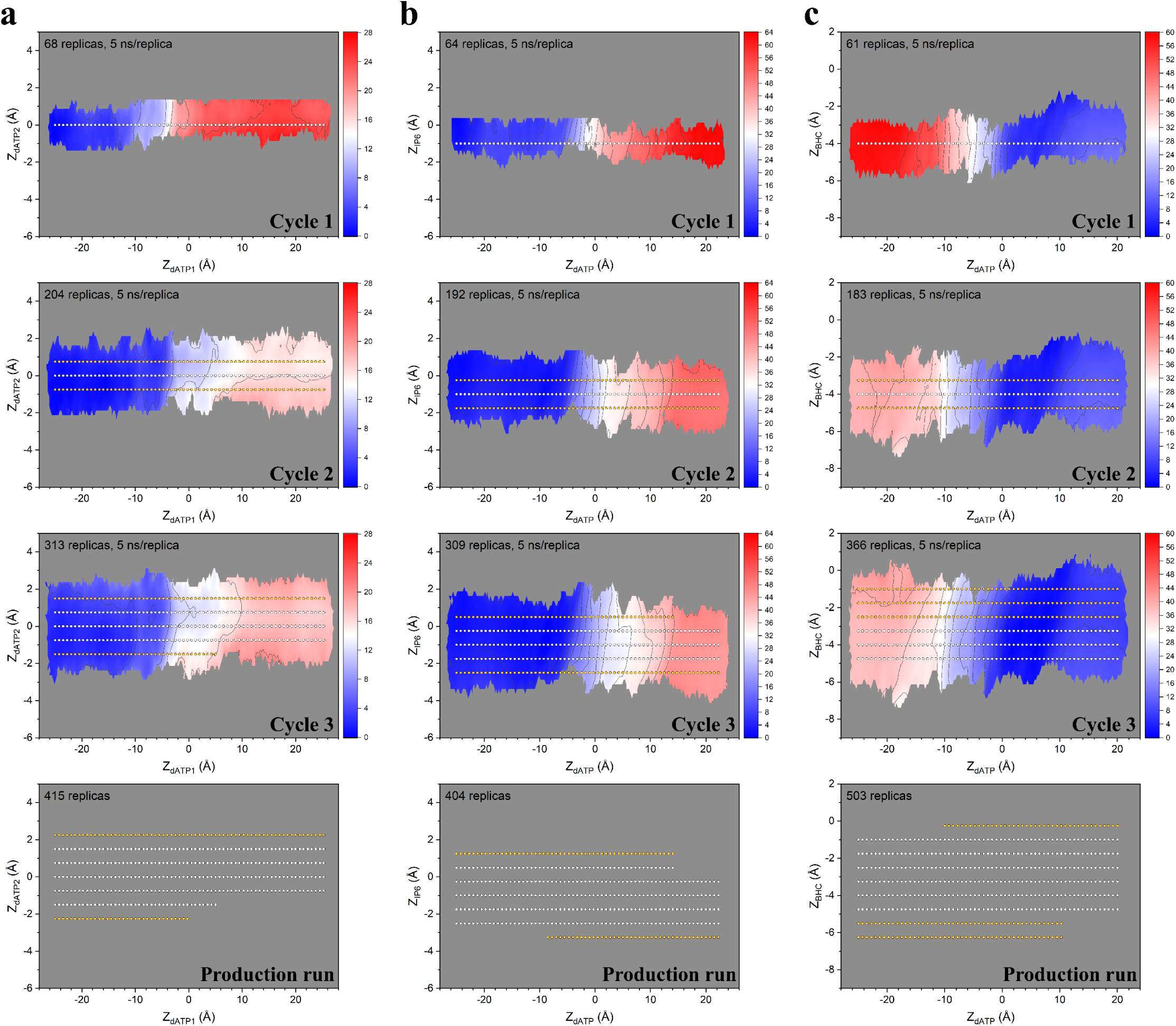
Setup of the 2D HREX/US simulations. (a) Generation of initial seeds for 2D HREX/US simulations of the 2 dATP model, (b) a dATP with IP_6_ and (c) BHC model, through 3 cycles of 5 ns HREX/US simulations. In each cycle, new US windows were generated to increase the sampling in 2D space. The initial conformations of the first cycles were extracted from SMD simulations pulling dATP through the hexamer central pore. After that, the initial conformations for new US windows (shown as yellow dots) were copied from the last frame of nearest previous US windows (shown as white dots). After 3 cycles, the generated seeds were used in 30 ns production runs.

**Figure S11.**
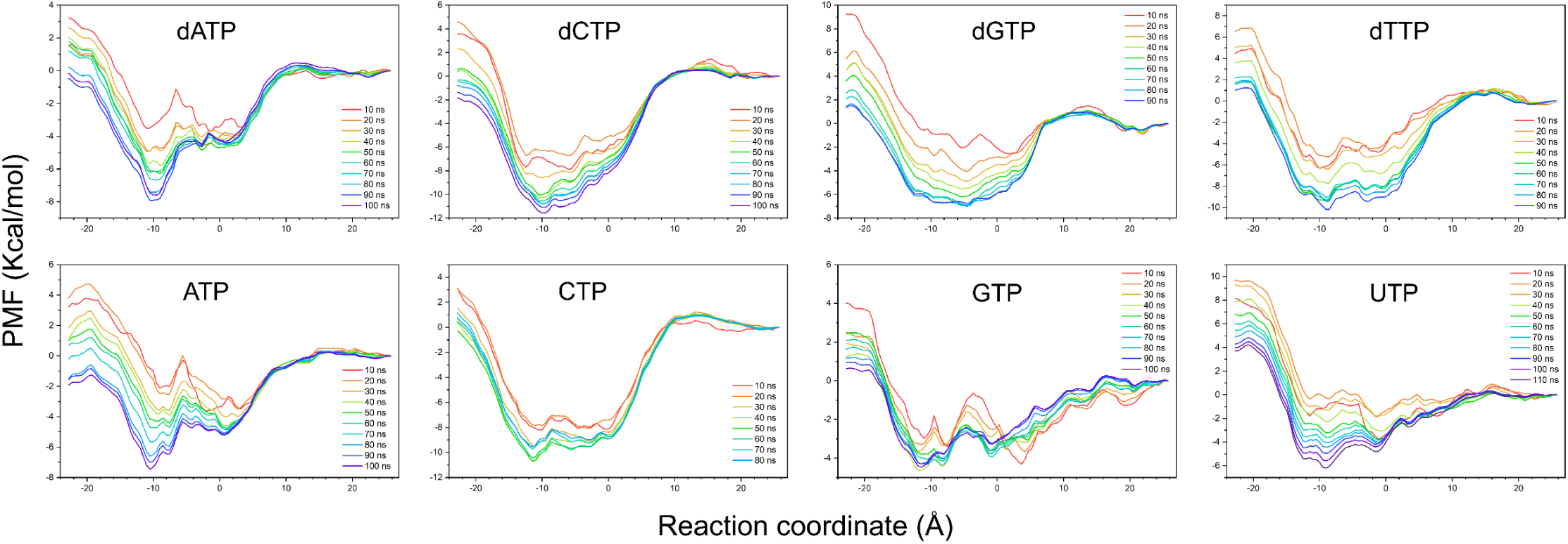
Convergence of the 1D HREX/US simulations. Sequential changes of PMF from every 10 ns HREX/US simulations are shown. After 10 ns, the PMF profiles were used to compute the 1D PMF for NTP translocation.

**Figure S12.**
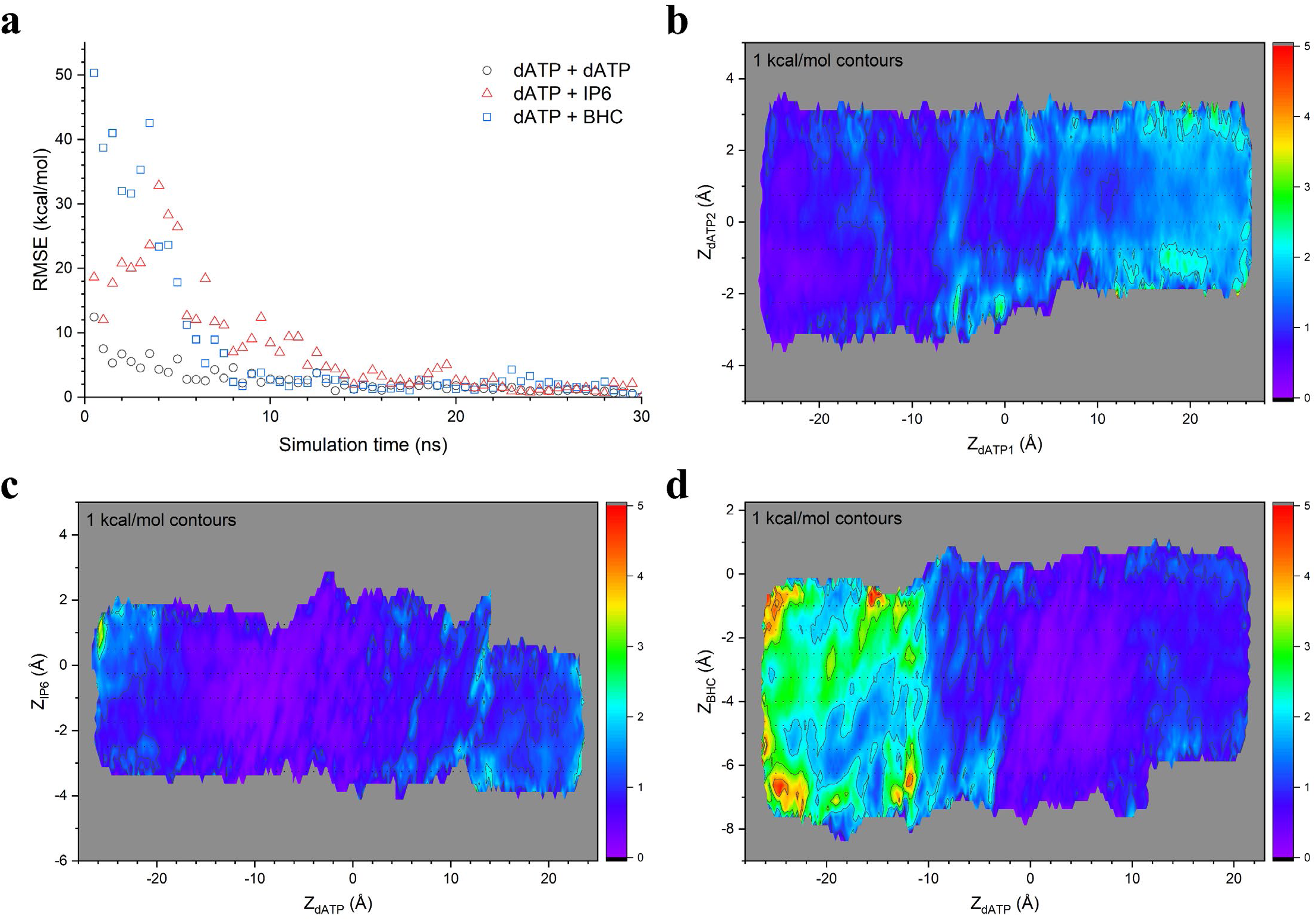
Convergence of the 2D HREX/US simulations. (a) Root mean squared errors of three 2D HREX/US simulations with 0.5 ns. Standard deviations of the 2D PMF surface of (b) the 2 dATP model, (c) dATP with IP_6_ and (d) dATP with BHC.

**Figure S13.**
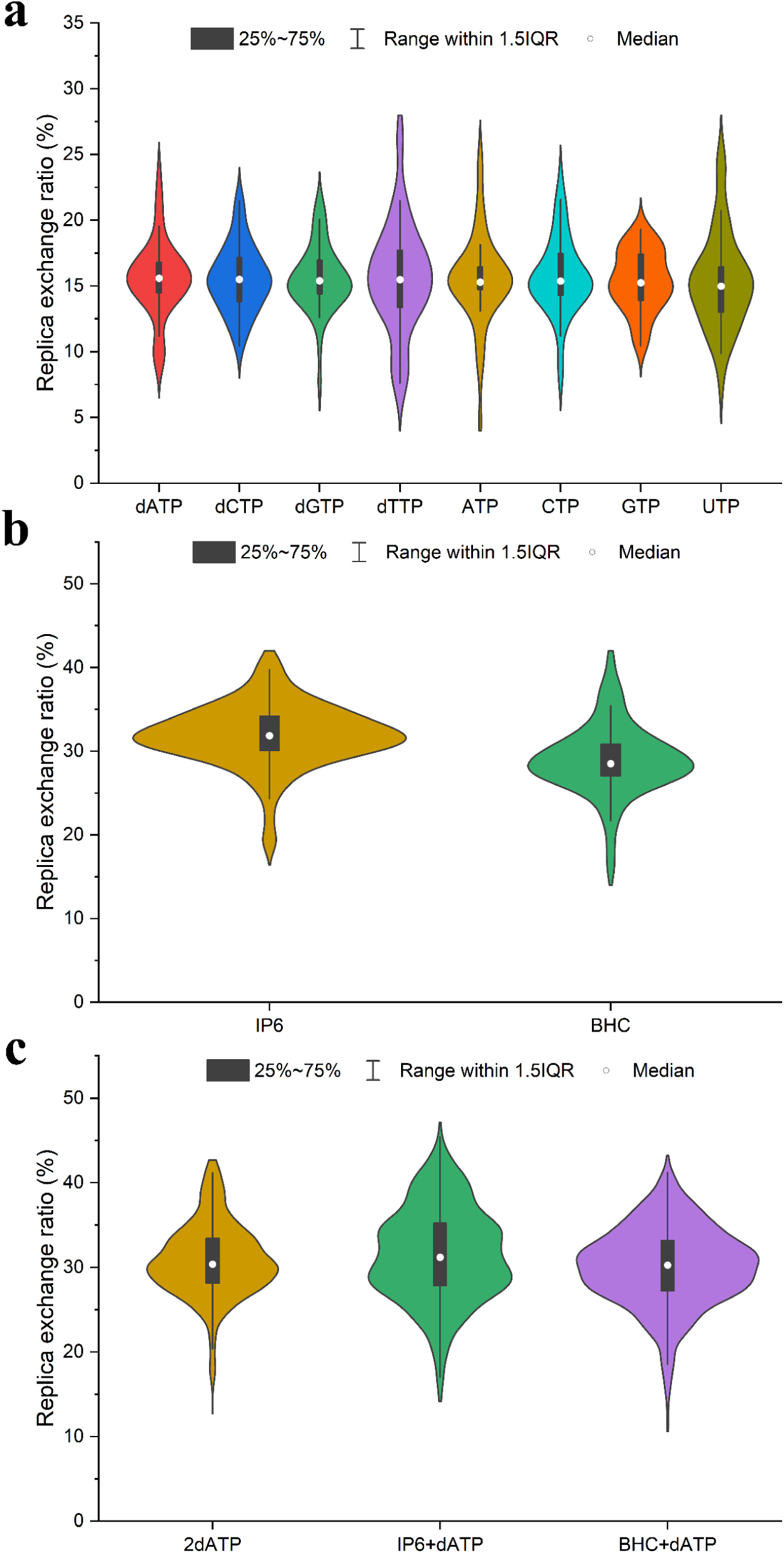
Exchange ratios for the HREX/US simulations performed in the present study. (a) 1D rNTP translocation simulation, (b) 1D IP_6_ and BHC translocation simulations and (c) the three 2D dATP translocation simulations.

**Figure S14.**
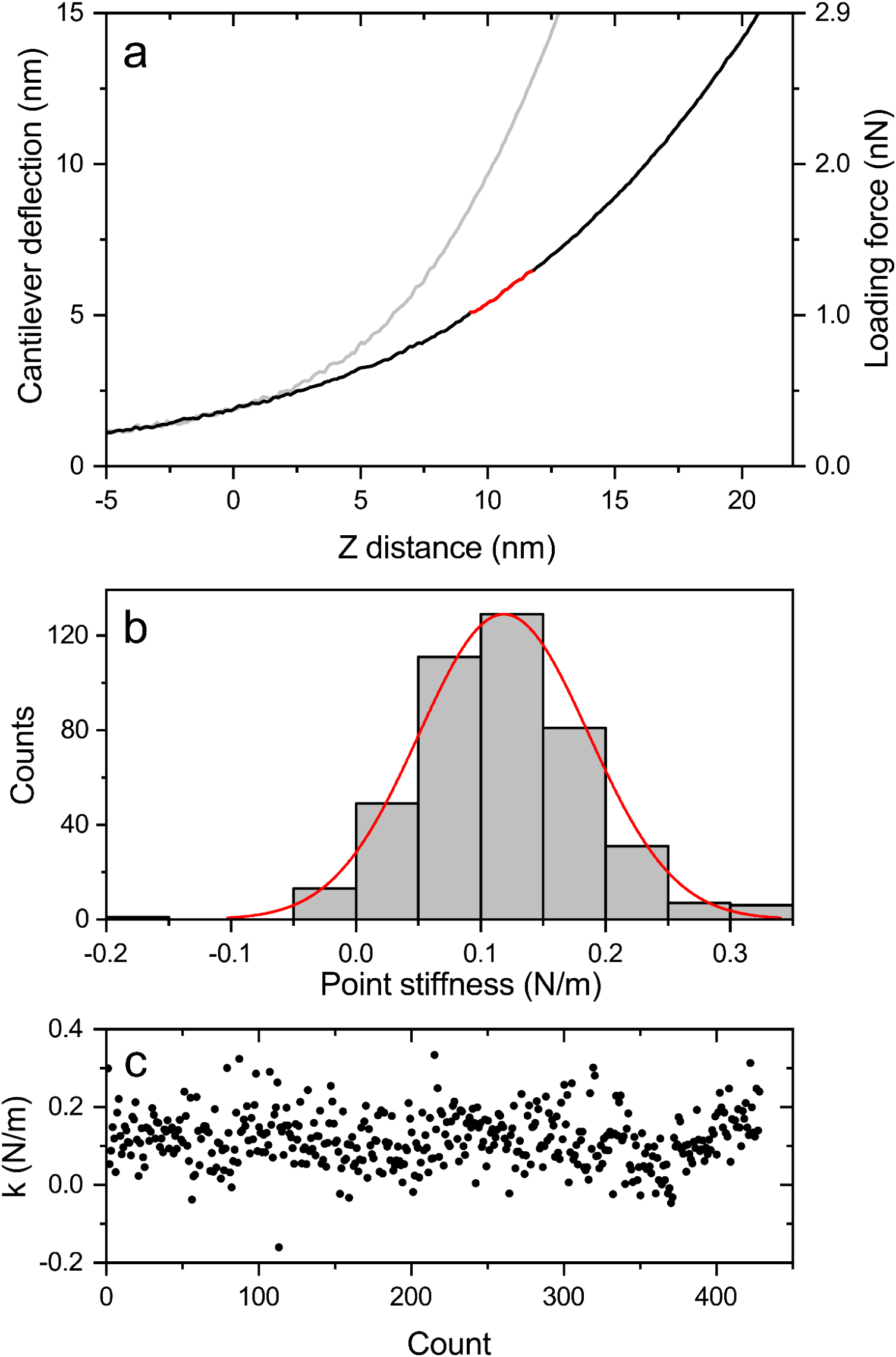
Measuring the point stiffness of the HIV-1 capsid by indentation-type experiments. (a) Histogram and Gaussian-fitted curves of the individual measured point stiffness values derived from the consecutive force-distance curves of a single core attached to a glass slide. (b) The individual measured point stiffness values obtained for the core shown in panel a, during a single experiment against the experiment number (count). The point stiffness measurements plot together with an analysis of the narrow distribution of the individual measured spring constants demonstrate that the core did not undergo a significant irreversible deformation during the indentation measurements. (c) Typical averaged force-distance curve of a core attached to an HMDS pretreated glass slide. For each experiment, ∼400 such curves were acquired and averaged. Stiffness values were calculated by fitting a linear function to the force curve region bounded by 3- and 4-nm indentation depths. The corresponding line fit is plotted in red.

**Table S1.**
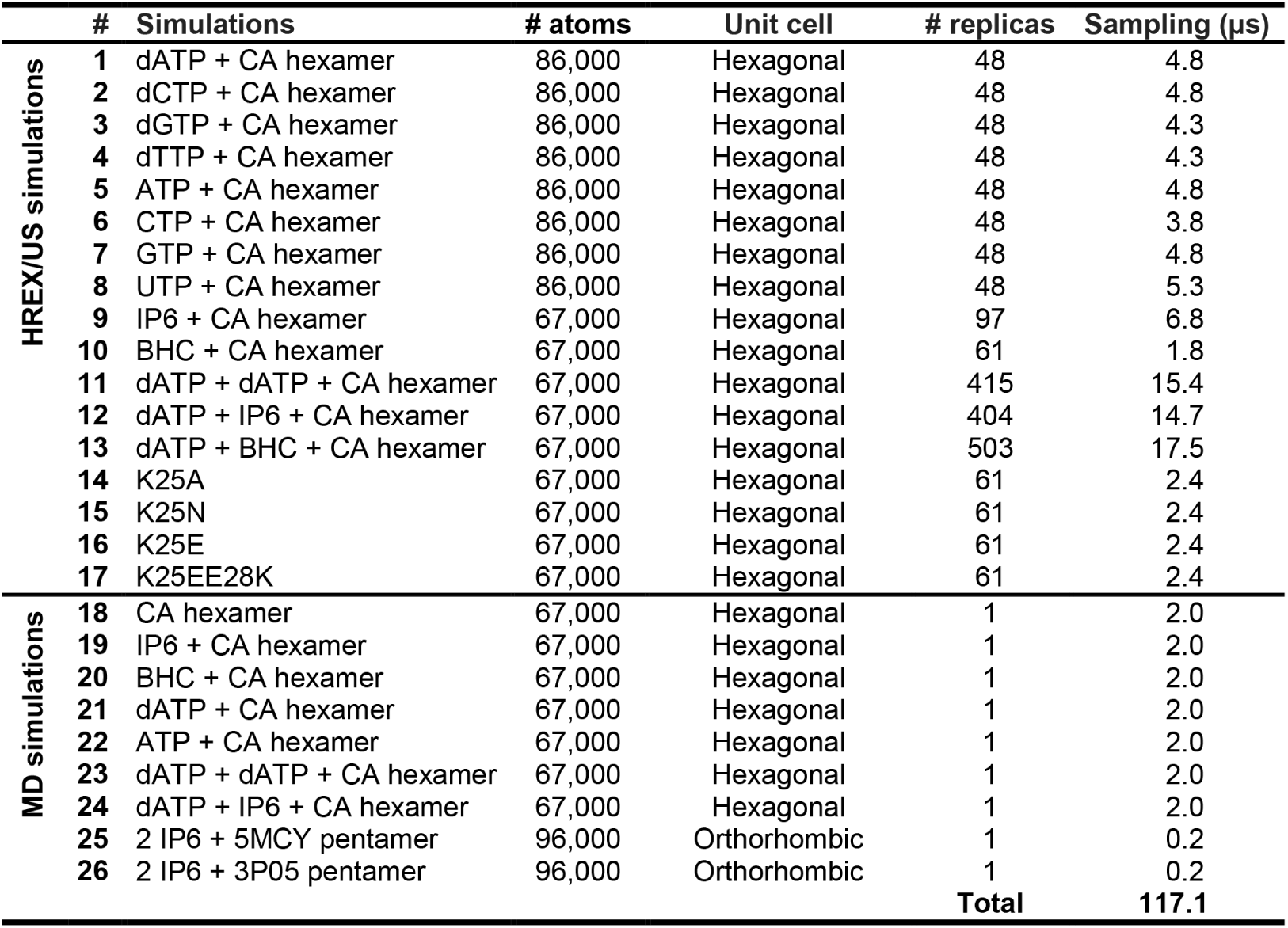
Simulations performed in the present study.

| | # | Simulations | # atoms | Unit cell | # replicas | Sampling (μs) |
| --- | --- | --- | --- | --- | --- | --- |
| HREX/US simulations | 1 | dATP + CA hexamer | 86,000 | Hexagonal | 48 | 4.8 |
|  | 2 | dCTP + CA hexamer | 86,000 | Hexagonal | 48 | 4.8 |
|  | 3 | dGTP + CA hexamer | 86,000 | Hexagonal | 48 | 4.3 |
|  | 4 | dTTP + CA hexamer | 86,000 | Hexagonal | 48 | 4.3 |
|  | 5 | ATP + CA hexamer | 86,000 | Hexagonal | 48 | 4.8 |
|  | 6 | CTP + CA hexamer | 86,000 | Hexagonal | 48 | 3.8 |
|  | 7 | GTP + CA hexamer | 86,000 | Hexagonal | 48 | 4.8 |
|  | 8 | UTP + CA hexamer | 86,000 | Hexagonal | 48 | 5.3 |
|  | 9 | IP6 + CA hexamer | 67,000 | Hexagonal | 97 | 6.8 |
|  | 10 | BHC + CA hexamer | 67,000 | Hexagonal | 61 | 1.8 |
|  | 11 | dATP + dATP + CA hexamer | 67,000 | Hexagonal | 415 | 15.4 |
|  | 12 | dATP + IP6 + CA hexamer | 67,000 | Hexagonal | 404 | 14.7 |
|  | 13 | dATP + BHC + CA hexamer | 67,000 | Hexagonal | 503 | 17.5 |
|  | 14 | K25A | 67,000 | Hexagonal | 61 | 2.4 |
|  | 15 | K25N | 67,000 | Hexagonal | 61 | 2.4 |
|  | 16 | K25E | 67,000 | Hexagonal | 61 | 2.4 |
|  | 17 | K25EE28K | 67,000 | Hexagonal | 61 | 2.4 |
| MD simulations | 18 | CA hexamer | 67,000 | Hexagonal | 1 | 2.0 |
|  | 19 | IP6 + CA hexamer | 67,000 | Hexagonal | 1 | 2.0 |
|  | 20 | BHC + CA hexamer | 67,000 | Hexagonal | 1 | 2.0 |
|  | 21 | dATP + CA hexamer | 67,000 | Hexagonal | 1 | 2.0 |
|  | 22 | ATP + CA hexamer | 67,000 | Hexagonal | 1 | 2.0 |
|  | 23 | dATP + dATP + CA hexamer | 67,000 | Hexagonal | 1 | 2.0 |
|  | 24 | dATP + IP6 + CA hexamer | 67,000 | Hexagonal | 1 | 2.0 |
|  | 25 | 2 IP6 + 5MCY pentamer | 96,000 | Orthorhombic | 1 | 0.2 |
|  | 26 | 2 IP6 + 3P05 pentamer | 96,000 | Orthorhombic | 1 | 0.2 |
|  |  |  |  |  | **Total** | **117.1** |

## Notes

### Competing Interest Statement

The authors have declared no competing interest.

